# Multi-omics Identification and Route-Specific Characterization of Metastasis-specific EMT Genes and Their Microenvironmental Interactions

**DOI:** 10.1101/2023.10.15.562367

**Authors:** Ki Tae Kim, Jae Eun Lee, Jae-Ho Cheong, In Cho, Yoon Young Choi

**Author notes:** **Correspondence: Ki Tae Kim, Ph.D.**, Department of Molecular Genetics & Dental Pharmacology, School of Dentistry, Seoul National University, Seoul, Korea, Zip: 08826, **Yoon Young Choi, M.D., Ph.D.**, Department of Surgery, Soonchunhyang Bucheon Hospital, Soonchunhyang University College of Medicine, 170 Jomaru-ro, Wonmi-gu, Bucheon-si, Gyeonggi-do, Republic of Korea, Zip: 14584. KT Kim and JE Lee are joint co-first authors.

## Abstract

**Background:** Gastric cancer (GC) constitute a significant cause of cancer-related mortality worldwide, with metastatic patterns including hematogenous, peritoneal, and ovarian routes. Although GC gene expression patterns have been extensively researched, the metastasis-specific gene expression landscape remains largely unexplored.

**Methods:** We undertook a whole transcriptome sequencing analysis of 66 paired primary and metastatic (hematogenous, peritoneal, or ovarian) GC tumors from 14 patients. Public databases including The Cancer Genome Atlas (TCGA) and Gene Expression Omnibus (GEO) was used for validation. Single-cell RNA sequencing (scRNA-seq) of four ascites from serosa positive GC patients and five primary tumors by layer (superficial and deep) were analyzed.

**Results:** Through differential expression analysis between paired primary and metastatic tumors by routes identified 122 unique metastasis-specific epithelial-mesenchymal transition (msEMT) genes. These genes demonstrated varying expression patterns depending on the metastatic route, suggesting route-specific molecular mechanisms in GC metastasis. High expression of msEMT genes in primary tumors was associated with more frequent *CDH1* mutations, the genomically stable subtype, and poor prognosis in TCGA GC cohort. This association was further corroborated by poor prognosis and high predictive performance for peritoneal/ovarian recurrence in two independent cohorts (GSE66229; n=300, GSE84437; n=433). scRNA-seq analysis of five primary tumors (GSE167297) and four independent ascites samples from GC patients revealed that msEMT genes were predominantly expressed in diverse fibroblast sub-populations, rather than cancer cells.

**Conclusions:** This study illuminates the route-specific mechanisms and underlines the significance of msEMT genes and cancer-associated fibroblasts in peritoneal metastasis of GC.

## Background

Gastric cancer (GC) is among the most prevalent malignancies worldwide, resulting in high cancer-related mortality. [1, 2] The primary cause of cancer-related deaths is distant metastasis, which in GC can be categorized into hematogenous, peritoneal, and specifically in females, ovarian metastasis. [3, 4] Among these, peritoneal metastasis is the predominant pattern in GC, and is linked with poor chemotherapy response and an ultimately unfavorable prognosis. [5]

Extensive research into the molecular characteristics of GC has improved our understanding and led to advancements in precision oncology. Notably, The Cancer Genome Atlas (TCGA) identified four molecular subtypes of GC: [6] microsatellite instability (MSI), Epstein-Barr Virus (EBV) related, chromosomal instability (CIN), and genomically stable (GS) subtype. Each of these presents unique histopathological, molecular, and clinical features. [6–10] Furthermore, these subtypes are associated with distinct metastatic patterns; hematogenous metastasis is associated with the CIN subtype, with increased copy number alterations compared to primary tumors, while peritoneal and ovarian metastasis are linked to the GS subtype, where chromosomal stability is maintained, yet *de novo* mutations arise. [11]

The patterns of gene expression in GC and their clinical implications have been extensively studied. [8, 12, 13] However, there remains a paucity of information on gene expression in metastatic GC and its corresponding biological and clinical implications, particularly in relation to metastatic patterns. To bridge this gap in knowledge, we conducted whole transcriptome sequencing analysis in paired primary and metastatic GCs. We found the route-specific mechanisms of GC metastasis and identified a unique gene set comprised of 122 metastasis-specific epithelial-mesenchymal transition (msEMT) genes. The expression of msEMT genes in primary tumors was related to poor prognosis and exhibited robust predictive power for peritoneal/ovarian metastasis, implying that GC might already possess molecular features conducive to peritoneal metastatic potential. Further, single-cell RNA (scRNA) sequencing analysis of the primary tumor, categorized by its layer, and ascites in patients with serosa positive GC revealed that msEMT genes predominantly originate from the microenvironment, specifically various fibroblasts, rather than cancer cells themselves.

## Methods

### Patients and tumor samples

Total 66 paired primary and metastatic tumors by their routes (38 primary, 9 of hematogenous, 6 of peritoneal, 13 of ovarian metastasis) were selected in 14 patients with GC and synchronous or metachronous metastasis. Formalin Fixed Paraffin Embedded (FFPE) tissues of tumors were collected. For single-cell RNA (scRNA) sequencing, intraperitoneal washing cytology for serosa positive GC before gastrectomy or ascites of patients with peritoneal metastasis were collected. Cytology was evaluated by pathologists and confirmed cytology positive or negative by the presence of malignant cells in ascites.

The patients had been treated following the gastric cancer treatment guidelines. [14] The clinical demographics of the enrolled patients and samples are described in Supplementary Tables 1 and 2. This study was approved by the Institutional Review Board (IRB) of Severance Hospital (4-2019-0188) and Soonchunhyang University Bucheon Hospital (2022-09-008), and informed consent was received for all patients.

### Whole Transcriptome sequencing (WTS)

For tissue sequencing, the FFPE of collected surgical specimens that were histologically confirmed as tumor samples were used. The histology of all samples was reviewed by a GC specialized pathologist, and RNA was obtained by macro-dissection of serial unstained sections from the tumor-enriched area. Subsequent WTS analysis to investigate gene expression patterns associated with gastric cancer metastasis. Multi-regional WTS was conducted for the primary tumors by the depth of tumors.

Total RNA was extracted from the FFPE tissue samples using a commercially available extraction kit (RNeasy FFPE kit, Qiagen, Germany), following the manufacturer’s protocol. The concentration and quality of the extracted RNA were assessed using an Agilent 2100 Bioanalyzer. Only RNA samples with satisfactory integrity, as indicated by an DV200 greater than or equal to 40%, were used for downstream applications. Library preparation and sequencing were performed by Macrogen, a reputable sequencing service provider. RNA libraries were prepared following the manufacturer’s instructions (TruSeq Stranded Total RNA Library Prep Kit, Illumine^TM^, CA, USA), were then sequenced on the Illumina NovaSeq6000 sequencing system (Illumina^TM^, CA, USA), generating paired-end reads with a read length of 101 bp. The raw paired-end sequencing data were aligned to the human reference genome (hg19) using the Spliced Transcripts Alignment to a Reference (STAR) aligner. [15] RNA sequencing analysis was performed using the bcbio-nextgen pipeline. [16] Gene expression quantification was carried out using the Salmon by tximport, [17, 18] which provided transcript-level quantifications that were subsequently aggregated to the gene level. Differential expression analysis was conducted using DESeq2, [19] with a false discovery rate (FDR) threshold of less than 0.05 or 0.01 to define significantly differentially expressed genes. To account for potential signal contamination due to the presence of normal cells in metastatic tumors, we used expression data from the GTEx database [20] and our internal data as a reference. The reference dataset consisted of normal liver, peritoneum, and ovary expression data. Genes highly or lower expressed in these normal tissues were considered potential contamination sources and their signals were adjusted accordingly in our differential gene expression analysis.

To further interpret our differential expression results, we performed Gene Set Enrichment Analysis (GSEA) on the lists of differentially expressed genes. Over-representation analysis was performed to identify biological pathways that were differentially expressed. The database for annotation were used Kyoto Encyclopedia of Genes and Genomes (KEGG), [21] PANTHER pathway [22] and Gene ontology of biological process (GO-BP) terms [23] to facilitate high-throughput gene functional analysis. The statistical significance was performed using a hypergeometric test. The significance threshold is less than 0.01.

### The Cancer Genome Atlas (TCGA) data analysis

We evaluated the biologic and genomic characteristics of the selected metastatic tumor specific gene set, we used TCGA of GC (STAD), including genomic data of 415 patients with GC. [6] Data (level 3) processed using an integrated annotation pipeline were downloaded from the National Cancer Institute Genomic Data Commons by TCGAbiolinks. [24] We performed unsupervised clustering using non-negative matrix factorization (NMF), by “*NMF*” package. [25] The optimal number of clusters was determined considering cophenetic value. Genomic characteristics of clusters determined by NMF were compared by “*maftools*”. [26]

To further explore the relationships between the selected genes, we conducted weighted gene co-expression network analysis (WGCNA) by “*WGCNA*” package. [27] After identifying modules of highly correlated genes, we selected a core module to identify NMF clusters and define a key gene set for msEMT genes.

### Gene Expression Omnibus data for primary tumors

The expression of selected msEMT genes was evaluated in two independent primary GC cohorts, ACRG (GSE66229, n=300) [8] and YCC (GSE84437, n=433). [13] Unsupervised clustering using Euclidean distance with the Ward.D method was used for each cohort, and the overall survival and disease free survival by the clusters were evaluated. Kaplan-Meier curves with log-rank test and cox-proportional hazard model were used and displayed with hazard ratio (HR) with its 95% confidence interval (CI). Age, sex, and TNM stage were used as adjustable variables for HR to identify the real prognostic effect of the msEMT genes.

### Machine learning algorithm to predict peritoneal/ovarian metastasis by msEMT genes

To assess the predictive performance of the msEMT genes for peritoneal or ovarian metastasis, we employed a variety of machine learning algorithms, including both classical machine learning techniques and state-of-the-art deep learning methods. Within the ensemble models group, we utilized Elastic Net regression (EN), Gradient Boosting Machine (GBM), and Random Forests (RF). These models combine multiple weak learners to form a strong predictive model. For kernel methods, we employed Support Vector Machine (SVM). This method is based on finding a hyperplane that optimally separates the data points into different classes. In the artificial neural networks category, we utilized Probabilistic Classifier Artificial Neural Networks (PCANN). This approach involves training a network of interconnected artificial neurons to learn complex patterns and make probabilistic classifications. By employing these diverse machine learning algorithms, we aimed to comprehensively evaluate the predictive capabilities of the msEMT genes for peritoneal and ovarian metastasis. For each method, we partitioned our dataset into a training set and a validation set using a 0.8:0.2 split with 10-fold repeated cross validation in each data set. Each model was trained on the training set and its performance was evaluated on the validation set. Hyperparameter tuning was carried out using a grid search or random search approach as appropriate for each model. To compare the performance of different models, major important prediction metrics were extracted, including Positive Predictive Value (PPV), Negative Predictive Value (NPV), Sensitivity, and Specificity. Additionally, Accuracy and Area Under the Receiver Operating Characteristic Curve (AUROC) were employed as evaluation metrics. To facilitate a comparison with the Generalized Linear Model (GLM)-based TNM stage and Lauren’s criteria, the performance comparison involved the application of DeLong’s test.

### Gastric cancer cell line data

The expression of GC cell lines was from SRP078289, [28] the expression of msEMT genes were compared by the origin of cell lines (primary, hematogenous, and ascites) using independent t-test and the difference was displayed as log2 fold changes.

### Single-cell RNA (scRNA) Sequencing

For scRNA-seq analysis, public data by layer of diffuse type of GC (superficial and deep layer of primary tumor) data in 5 patients was obtained from GSE167297, [29] and internal data from ascites of 4 patients with serosa positive GC. We employed the Chromium Single Cell 3’ Protocol (10x Genomics) to profile 3’ digital gene expression in 500 - 10,000 individual cells per sample. Single cells, reagents, and a single Gel Bead encapsulating barcoded oligonucleotides were co-encapsulated into nanoliter-scale Gel Beads in Emulsion (GEMs) using the Next GEM technology. The single-cell 3’ gene expression (GEX) and feature barcode libraries were then generated following the manufacturer’s protocol. The library was prepared using the Chromium Next GEM Single Cell 3’ RNA library kit v3.1 (10x Genomics). The prepared libraries were sequenced on the Illumina sequencing system, with paired-end sequencing performed for Read 1 and Read 2 from both ends of the fragment. The raw sequencing data were demultiplexed and converted to FASTQ format using bcl2fastq software v2.20.0 (Illumina).

Each sample was loaded with more than 9,000 cells as the target output. The scRNA-Seq data were demultiplexed, aligned to the human genome (hg38), and UMI-collapsed using the CellRanger toolkit (version 6.0.1) provided by 10x Genomics with default parameters. The “mkfastq” module in CellRanger demultiplexes the raw base call (BCL) files generated by Illumina sequencers into FASTQ files. The total number of bases, reads, GC (%), Q20 (%), and Q30 (%) were calculated for four samples. The mean number of reads per cell was 32,324, 39,989, 46,444, and 32,607 in ascites S1, S2, S3, and S4, respectively. The UMI Q30 bases were 97.8%, 97.7%, 97.9%, and 97.5% in ascites S1, S2, S3, and S4, respectively. (Supplementary Table 3)

### Data preprocessing, batch correction and doublet elimination of scRNA-seq

Gene expression data were processed using the Seurat, v.4.9.9. R package. [30] We filtered out genes that were not expressed in at least 10 cells and excluded cells that had fewer than 200 detected genes. For feature filtering, cells with fewer than 2,500 counts and less than 5% mitochondrial content were removed. Gene expression levels were represented as the fraction of UMI counts relative to the total UMI counts in each cell and were normalized using size factor normalization, aiming for 10,000 counts per cell (TP10K). To correct batch effects in the multiple single-cell data, we employed the Seurat [30] and Liger [31] software. Seurat employs anchor-based integration, which aligns the datasets by identifying similar cell populations across batches and harmonizing their expression profiles using “FindIntegrationAnchors” and “IntegrateData” functions. Liger uses a network-based approach that constructs a shared nearest-neighbor graph to identify similar cells across batches and align their gene expression patterns. Finally, to remove computationally predicted doublet cells, we utilized the DoubletFinder. [32]

### Dimensional Reduction and Clustering of scRNA-seq

We employed dimensionality reduction techniques on the gene expression data using a subset of 1000 highly variable genes. The selection of variable genes was based on their dispersion of binned variance to mean expression ratios, accomplished using the “FindVariableGenes” function in the Seurat package. Additionally, we filtered out genes associated with the cell cycle, ribosomal proteins, and mitochondria. To reduce the dimensionality of the data, we performed principal component analysis (PCA) and retained the top 50 PCA components. The selection of the number of components was determined based on the standard deviations of the principal components, observing a plateau region on an elbow plot. We then utilized graph-based clustering on the PCA-reduced data, employing the Louvain Method after constructing a shared nearest neighbor graph. The resulting clusters were visualized using t-distributed stochastic neighbor embedding (t-SNE) and Uniform Manifold Approximation and Projection (UMAP) techniques to generate a 2D map. For sub-clustering in each cell type, we applied the same procedure of identifying differentially expressed genes within the analyzed cluster, followed by dimensionality reduction and clustering on the restricted subset of data, typically focused on a specific initial cluster.

### Cell type annotation and composition for scRNA-seq

Cell types with functional identities were assigned to cell clusters based on differential expression signatures derived from statistics of log likelihood ratio test. To define the cell clusters, we performed principal component analysis on the most variable genes between cells, followed by Louvain and Leiden graph-based clustering. We extracted significantly differentially expressed genes in each of the original raw clusters from the analyzed samples. For elaborate mapping between clusters and cell type, we used two algorithms with Single R [33] for robust immune cell mapping and scATOMIC [34] for adjusting immune and cancer cell signals. Finally, we confirmed cell types using manual curation based on the consensus of three researchers. When assigning the molecular characteristics for sub-cluster identity, we used top signature genes of up to most differentially expressed between each sub-cluster and others.

### Differential expression analysis (DEA) in scRNA-seq

For identifying remarkable markers in each cluster, we applied the Wilcoxon Rank-Sum Test using the “FindMarkers” function of Seurat package) to find genes that had significantly different RNA-seq TP10K expression when compared to the remaining clusters. The selection criteria for single cell clustering were more than two-fold change between each cluster and others. For differential expression analysis between fibroblast, we used a generalized linear model treating for comparison between metastatic ascites and primary GC by DESeq2.

The threshold of log2 fold change was more than four between two groups and all the false discovery rate (FDR) threshold was less than 0.01 to define significantly differentially expressed genes.

### Constructing trajectory inference for scRNA-seq

We employed Monocle 3 [35] to perform trajectory analysis on a scRNA-seq dataset. The dataset consisted of analyzed single cells and was preprocessed to remove batch effects, normalize gene expression, and identify highly variable genes. We utilized Monocle 3’s functionality to reduce the dimensionality of the dataset using principal component analysis (PCA) with Uniform Manifold Approximation and Projection (UMAP) for identifying the most informative genes driving cellular transitions. The dimensionality-reduced data was then fed into a pseudotime analysis algorithm to infer the temporal dynamics of the cellular development. Pseudotime ordering of cells was performed to uncover the continuous and nonlinear progression of cell states or cell differentiation using “learn_graph” and “order_cells” functions. The algorithm constructed to optical starting cell types with a minimum spanning tree or computed diffusion-like distances between cells to infer trajectories, branching events, and even loops. All trajectory analyses were performed individually for each dataset. The resulting values and distribution of pseudotime were interpreted as the cell’s relative progress along the developmental trajectory. We analyzed the pseudotime-dependent expression of genes to identify cellular progress that might drive the developmental process.

### Cellular communication analysis across cell types in scRNA-seq

To identify cell-cell communication events within the scRNA-seq data, we employed the CellChat toolkit, [36] a computational framework designed specifically for the analysis of cell-cell communication networks in scRNA-seq data. First, we utilized CellChat’s built-in algorithms to identify potential ligand-receptor interactions present in the dataset. This analysis involved examining the expression patterns of ligand and receptor genes across the different cell populations. Next, we calculated communication probability score, a metric used to quantify the strength of cell-cell communication. These scores were computed based on the expression levels of the identified ligand-receptor pairs, employing the statistical models and algorithms provided by the CellChat toolkit. For network analysis, we constructed a cell-cell communication network, where each node represented an individual cell type or cell population, and the edges represented the communication interactions between them. Within these pathway-specific communication networks, we calculated various centrality measures such as degree centrality, betweenness centrality, and closeness centrality for each cell type.

### Statistical analysis

The data are presented as mean ± standard deviation and compared using the student’s *t*-test or Mann–Whitney U test. For survival analysis, overall survival was defined as death from a GC diagnosis, and it was generated by Kaplan–Meier curves with a log-rank test. For multivariable analysis, the Cox proportional hazard model was used and displayed using the hazard ratio (HR) and its 95% confidence interval (CI). R version 4.2.2 (R Foundation for Statistical Computing, Vienna, Austria) was used for all statistical analyses. For multiple hypothesis correction of statistical tests were performed using Benjamin-Hochberg’s method.

## Results

### Identification of Metastatic GC Specific Genes

To identify metastatic GC specific genes, we performed a differential expression analysis, comparing genes expressed more in primary tumors (n=38) against normal gastric tissues (n=4). Then we analyzed the expressions within matched primary and metastatic tumors, considering the distinct metastatic routes (hematogenous, peritoneal, and ovarian, Figure 1A). Genes with up-regulated in metastatic tumors, yet their expression was ≥ 80% of the quantile in the corresponding normal tissue (e.g., liver, peritoneum, ovary, respectively) were excluded to remove the impact of normal contamination. We also excluded down-regulated genes with an expression ≤ 20% of the quantile (Figure 1A and Supplementary Figure 1).

**Figure 1.**
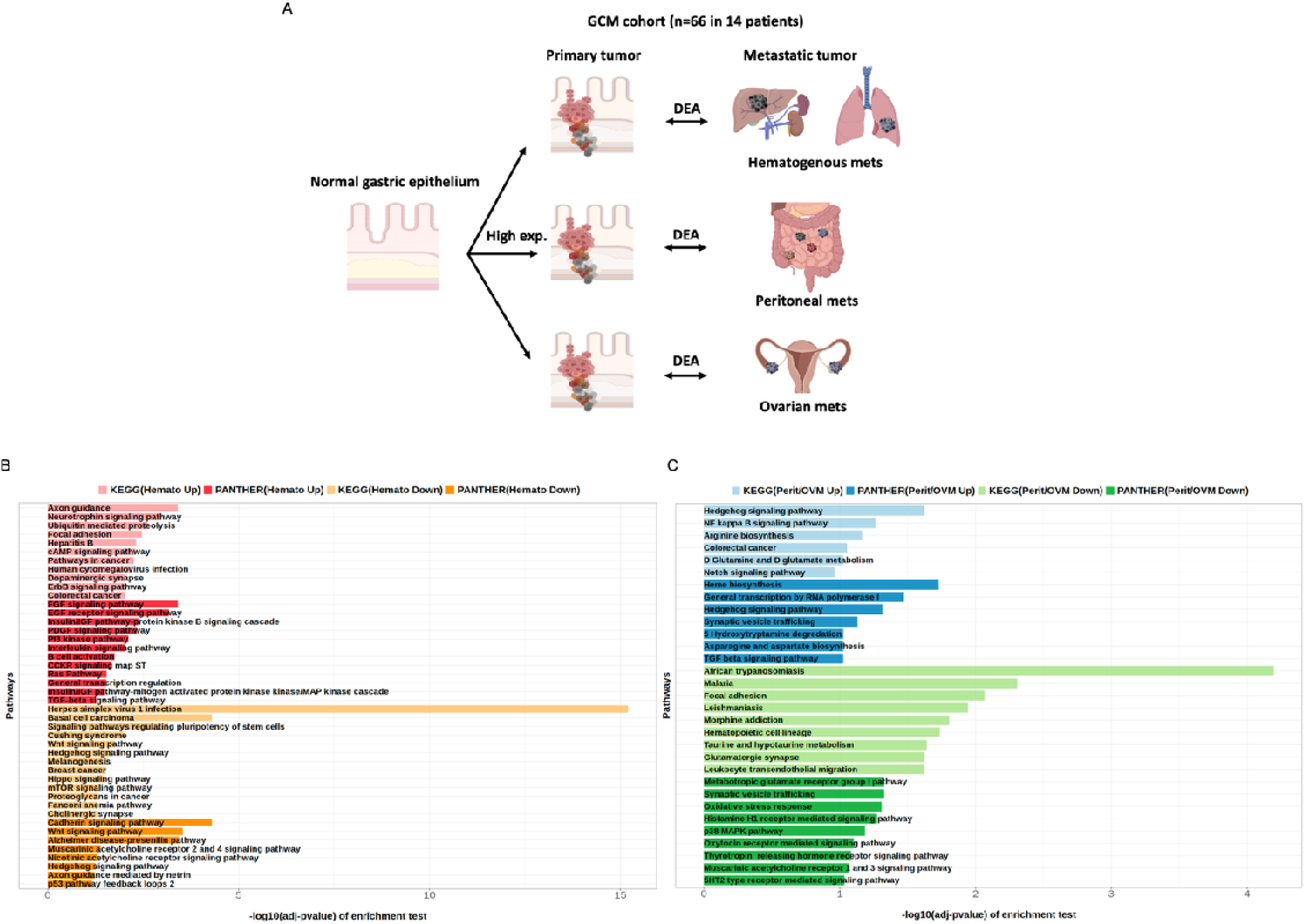
Scheme of differential expression analysis conducted in this study and over-representation analysis of metastatic gastric cancer-specific genes. A) Initial selection involved genes highly expressed in primary gastric cancer (n=38) compared to normal gastric tissues (n=4). Next, expression within matched primary and metastatic tumors, considering metastatic routes (hematogenous, peritoneal, and ovarian metastasis), was compared. Through this analysis, we identified metastatic gastric cancer-specific genes. B) Over-representation analysis of metastatic gastric cancer-specific genes for hematogenous metastasis. C) Over-representation analysis of metastatic gastric cancer-specific genes for peritoneal/ovarian metastasis. DEA; differential expression analysis, OVM; ovarian metastasis

A subsequent differential expression analysis between metastatic tumors and other tumors, respective to their metastatic routes in the overall sample pool, identified a total of 949 genes. Over-representation analysis and gene set enrichment analysis (GSEA) for hematogenous and peritoneal/ovarian metastasis (In our previous study, ovarian metastasis has similar characteristics with peritoneal metastasis in GC) [11] revealed up-regulation of Receptor Tyrosine Kinase (RTK) related signals, such as Fibroblast Growth Factor (FGF), Epidermal Growth Factor receptor (EGFR), and Platelet-Derived Growth Factor (PDGF) signaling pathways in hematogenous metastasis. Interestingly, Epithelial-Mesenchymal Transition (EMT) associated pathways (Wnt, Hedgehog, Hippo signaling) were down-regulated in this route. Conversely, the Hedgehog pathway, a representative EMT signaling pathway, was amplified in peritoneal/ovarian metastatic tumors. This suggests a significant divergence in pathway enrichment, contingent on the metastatic routes in GC (Figure 1B-C and Supplementary Figures 2).

### Expression of Metastatic GC Specific Genes in Primary GC

To investigate the clinical and biological implications of the identified metastatic GC-specific genes, we utilized The Cancer Genome Atlas (TCGA), a comprehensive primary tumor database. With our selection of genes, we found that GC could be stratified into two distinct subtypes using Non-negative Matrix Factorization (NMF_G1 and NMF_G2, see Supplementary Figure 3A and Figure 2A). The NMF_G2 group was strongly associated with the GS subtype, less so with MSI or EBV subtypes (Supplementary Figure 3B). Importantly, higher expression of these genes (NMF_G2 group) was associated with a worse prognosis, both overall and within each molecular subtype as suggested by TCGA (Figure 2B, Supplementary Figure 3C). The *CDH1* somatic mutation was positively associated with NMF_G2 (Figure 2C and Supplementary Figure 3D). We reclassified the molecular subtypes of the TCGA cohort into msEMT (metastatic GC specific EMT; NMF_G2 group), MSI, EBV, and non-msEMT types. The prognosis stratified by these revised subtypes demonstrated superior separation compared to the original subtypes (Figure 2D). We identified four modules using Weighted Gene Co-expression Network Analysis (WGCNA) (Supplementary Figure 4). A specific module of 122 genes demonstrated distinctive characteristics between NMF_G1 and NMF_G2. Over-representation analysis revealed that these genes were associated with EMT-related signals such as Hedgehog, Hippo, Wnt, and cadherin signaling pathways (Figure 2E), leading us to name this gene-set as metastasis-specific EMT (msEMT) genes.

**Figure 2.**
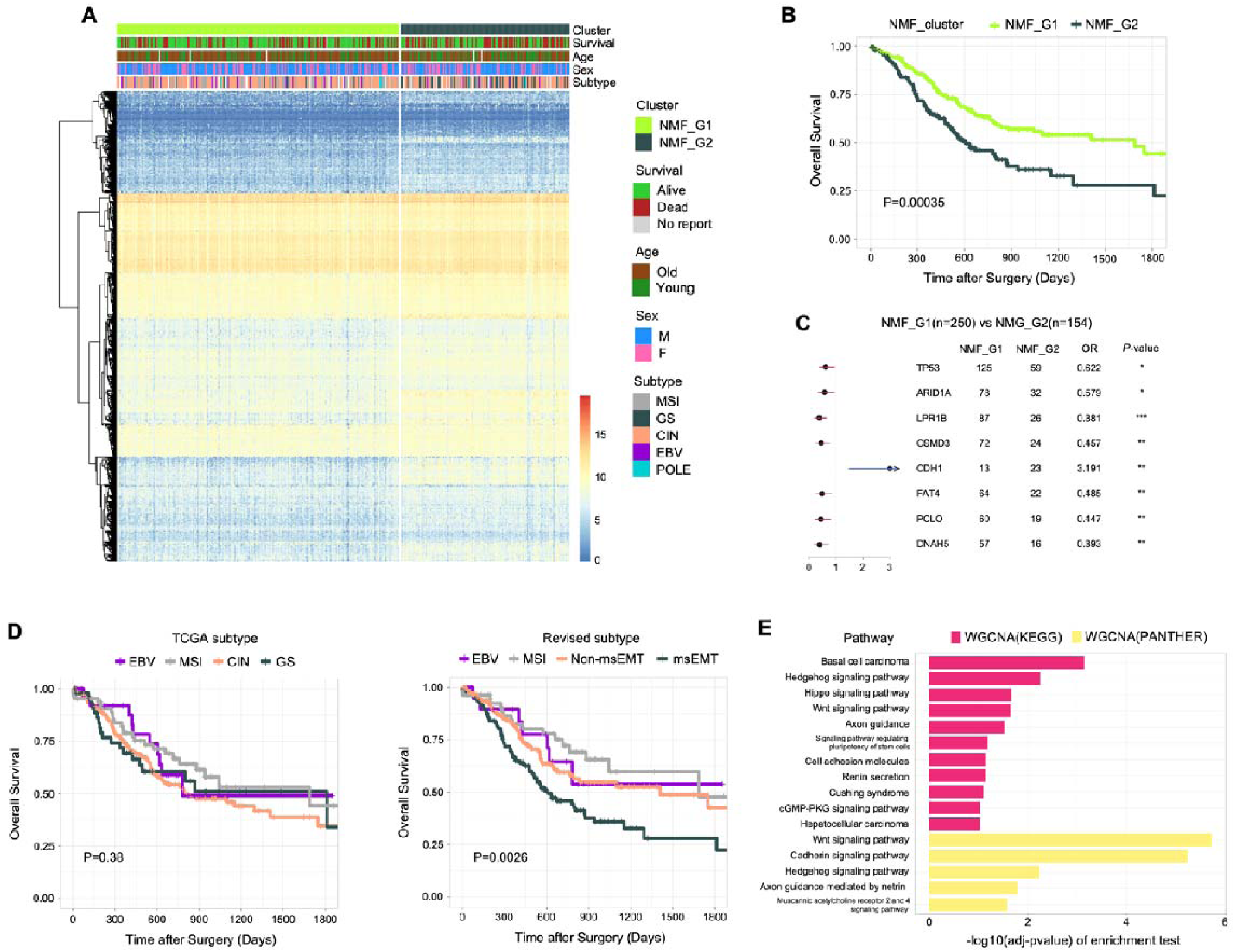
Clinical and biological characteristics of metastatic gastric cancer-specific genes in the cancer genome atlas (TCGA) cohort. A) Heatmap of metastatic gastric cancer-specific genes and their clinical characteristics. Non-negative matrix factorization (NMF) classified gastric cancer into two subtypes: NMF_G1 and NMF_G2. B) Kaplan-Meier curves of overall survival for NMF_G1 and NMF_G2. NMF_G2, which highly expresses metastatic gastric cancer-specific genes, demonstrated a poorer prognosis than NMF_G1. C) Comparison of the frequency of somatic mutations between NMF_G1 and NMF_G2. CDH1 mutations were frequently observed in NMF_G2. D) Kaplan-Meier curves of overall survival by molecular subtypes proposed by TCGA and revised subtypes, including metastatic gastric cancer-specific EMT type (NMF_G2). Prognosis was well stratified by revised subtypes compared to original subtypes. E) Over-representation analysis of the final selection of 122 metastasis-specific EMT (msEMT) genes. EMT-related signals such as Hedgehog, Hippo, Wnt, and cadherin signaling pathways were observed.

Comparing this set with well-established gene-sets in GC biology [8, 37–41] and consensus cancer driver genes, [42] most genes within the msEMT genes set were novel, suggesting the unique nature of this gene-set (Supplementary Tables 4). Interestingly, 103 of these genes were selected due to their down-regulation in hematogenous metastatic tumors compared to primary tumors. Upon re-evaluating the expression of msEMT genes in peritoneal/ovarian metastatic tumors and primary tumors, we found that most of them were highly expressed in the metastatic tumors (Supplementary Tables 5).

### Prognostic Impact of msEMT gene expression and its association with Peritoneal/Ovarian Metastasis

To substantiate the clinical implications of msEMT genes in GC, we assessed their expression in two independent primary GC cohorts: the ACRG (n=300) and the YCC (n=433) cohorts. [8, 13] The expression of msEMT genes enabled the classification of each cohort into two distinct subtypes (Figure 3A and B). Notably, the group with high expression, designated as the msEMT subtype, was associated with a worse prognosis (log rank-*p* = 0.0018, 0.011, respectively, Figure 3 C and D). This association held even after adjusting for clinical factors such as patients’ age, sex, and TNM stages, indicating a poorer prognosis in both cohorts (overall survival with adjusted HR: 1.66 [95% CI: 1.18-2.34], *p*=0.0035, and adjusted HR: 1.50 [95% CI: 1.13-1.98], *p*=0.0049, respectively).

**Figure 3.**
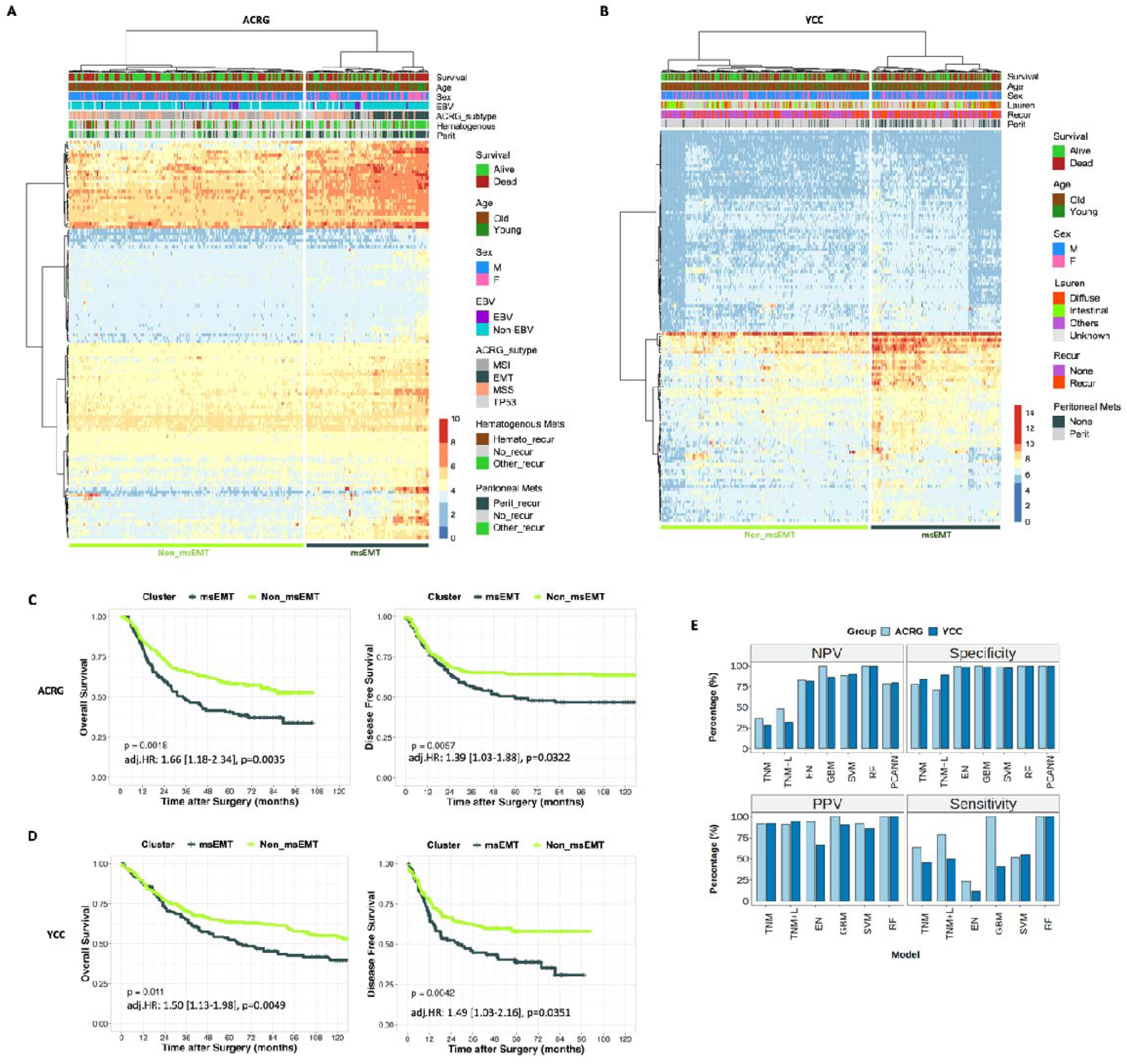
Clinicopathologic characteristics of metastasis-specific EMT (msEMT) genes in the ACRG (n=300) and YCC (n=433) cohorts. A) Heatmap by msEMT genes showed that high expression of msEMT genes was related to the frequent EMT subtype by ACRG and peritoneal but not hematogenous metastasis in the ACRG cohort. B) Heatmap by msEMT genes showed that high expression of msEMT genes was related to frequent diffuse histology and peritoneal recurrence in the YCC cohort. Kaplan-Meier curves for overall survival (OS) and disease-free survival (DFS) by the expression of msEMT genes C) in the ACRG cohort and D) in the YCC cohort. E) Performance in predicting peritoneal/ovarian metastasis with the expression of msEMT genes by various machine learning algorithms. Overall, their performance was better compared to the TNM stage or TNM + Lauren classification (logistic regression model), especially in terms of negative predictive value and specificity. TNM; tumor node metastasis, TNM+L; TNM + Lauren classification, EN; Elastic net, GBM; Gradient Boosting Machine, RF; random forest, SVM; support vector machine, PCANN; probabilistic classifier artificial neural networks.

Moreover, the msEMT subtype of primary tumors exhibited a stronger correlation with peritoneal/ovarian recurrence than with hematogenous metastasis (Figure 3A and B). We assessed the predictive power of msEMT genes for peritoneal/ovarian metastasis using various machine learning algorithms. The msEMT genes in the primary tumors demonstrated a high predictive accuracy (>0.80), and notably, they exhibited a high negative predictive value and specificity when compared to models that incorporated TNM stage or a combination of TNM and Lauren classification (Figure 3E, Supplementary Table 6). The models that combined TNM and Lauren classification also showed high predictive performance for peritoneal/ovarian metastasis. These findings suggest that msEMT genes play a pivotal role in peritoneal/ovarian metastasis, and primary GC already possesses the potential for such metastasis.

### Tracing the origin of msEMT gene expression

To understand the origin of the unique msEMT gene expression in GC, we first compared the expression of msEMT genes across GC cell lines, comprised of monoclonal cancer cells, based on their origin (primary, n=6; liver metastasis, n=3; ascites, n=10; Supplementary Table 7). Notably, no gene showed significantly higher expression in GC cells derived from malignant ascites compared to those from primary tumors, implying that msEMT signals might not originate from cancer cells themselves (Supplementary Table 8).

To further investigate this, we performed single-cell RNA sequencing (scRNA-seq) analysis on five primary tumors stratified by layer (superficial and deep layer, GSE167297) [29] and four ascites samples from patients with serosa-positive GC (internal data, including two samples with cytology-negative and two with peritoneal metastasis, Figure 4A). Our findings revealed that msEMT genes were not predominantly expressed in cancer cells. Instead, we observed their primary expression in endothelial cells within the superficial layer of the primary tumor, in endothelial cells and fibroblasts within the deep layers of the primary tumor, and in fibroblasts within the ascites (Figure 4 B-D). These findings suggest that msEMT signals may arise from the tumor microenvironment relating to cancer-associated fibroblasts (CAFs) in the late stage of primary tumor (deep layer) and peritoneal metastasis rather than cancer cells.

**Figure 4.**
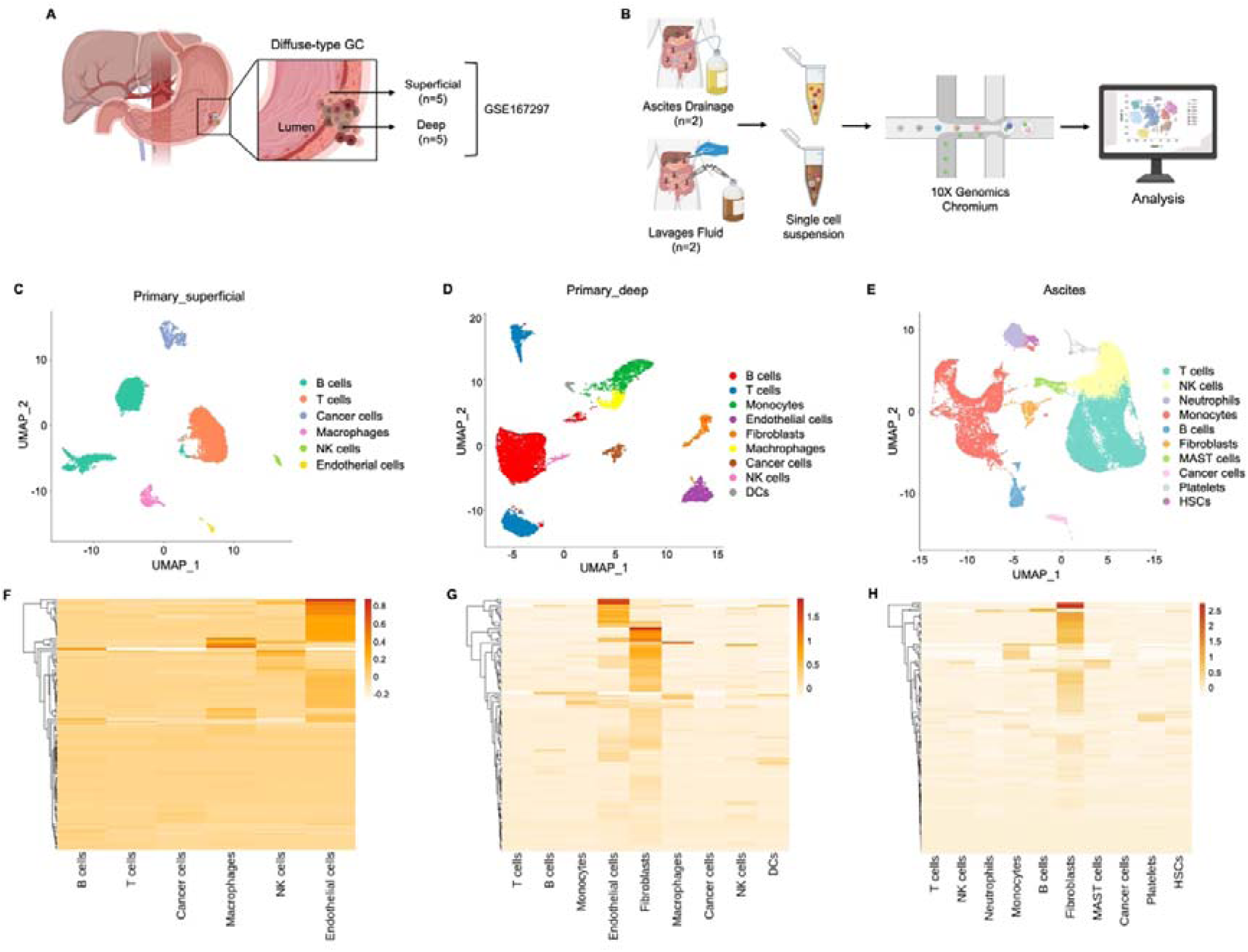
Single-cell RNA-seq profiles of msEMT genes in primary gastric cancer by layer (superficial and deep layer, n=5) and ascites (n=4). A) Schematic of the experimental design of this study for single-cell RNA-seq. (a) Public data (GSE167297), which includes single-cell RNA-seq in primary gastric cancer by layer and (b) ascites and peritoneal lavage fluid, was used for scRNA-seq analysis. Uniform Manifold Approximation and Projection (UMAP) (above) and the expression of msEMT genes in the main cell types (below) of B) the superficial layer of the primary tumor, C) the deep layer of the primary tumor, and D) ascites. The expression of msEMT genes was mainly observed in endothelial cells in the superficial layer, fibroblasts and endothelial cells in the deep layer, and fibroblasts in the ascites but were rarely expressed in cancer cells.

To further understand the expression patterns of msEMT genes within specific fibroblast populations, we explore the landscape and dynamic changes in fibroblasts in the primary tumor and ascites in detail. In the deep layer of the primary tumor, cell lineage trajectory of fibroblasts showed that C1 fibroblasts (*ACTA2+/S100A4+*), characterized as classical myofibroblastic CAFs (myo-CAFs), and C0 fibroblasts (*CXCL12+*), referred to as inflammatory CAFs (i-CAFs), were at the origin of the pseudotime trajectory. Meanwhile, C2 fibroblasts (*PDPN+/FN1+*), CAFs were associated with cancer migration and EMT, [43, 44] were at the end of the pseudotime trajectory (Figure 5A). The msEMT genes were primarily yet differentially expressed across C0, C1, and C2 fibroblasts. (See Figure 5A, right) In the ascites, C1 fibroblasts (*VEGFA+*) and C3 fibroblasts (*CXCL12+*), referred to as vascular and inflammatory CAFs (v- and i-CAFs), were at the origin of the pseudotime trajectory, with C5 fibroblasts (*ACTA2+/S100A4+*) characterized as myo-CAFs, in the middle, and C6 (*TNFRSF12A+*) and C7 (*S100A4+/FN1+*) fibroblasts, which may represent mature CAFs associated with CAF proliferation [45] and cancer cell migration, [46] at the end of the pseudotime trajectory (Figure 5B). The msEMT genes were mainly expressed across C1, C3, C6, and C7, but were rarely expressed in C5 fibroblasts (myo-CAFs). (See Figure 5B, right) This observation indicates that msEMT genes is associated with the diversity and specificity within the CAF population, potentially playing a role in modulating the fibroblast cellular landscape during the process of GC peritoneal metastasis.

**Figure 5.**
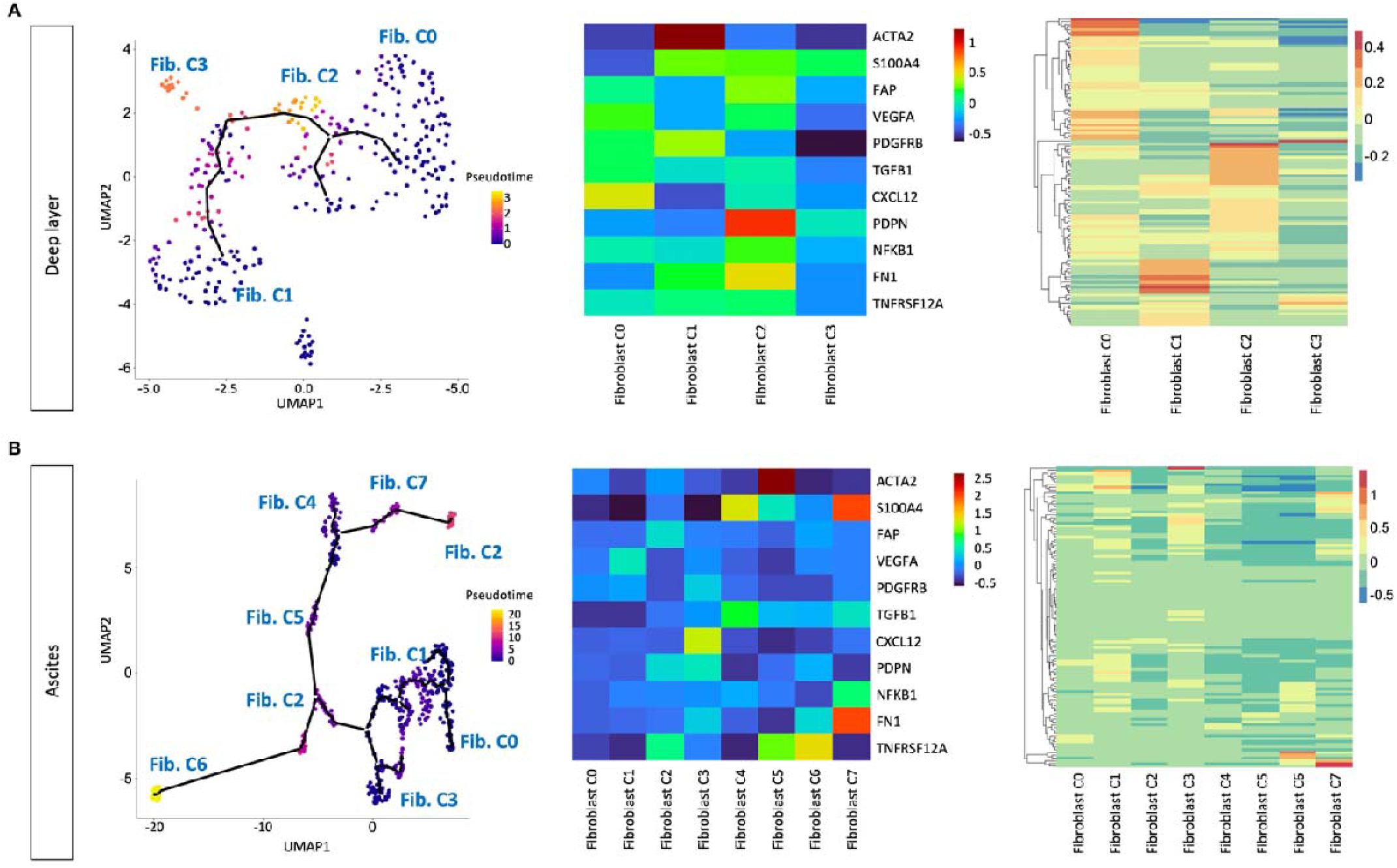
Diversity and dynamics of fibroblasts and the expression of msEMT genes in various sub-populations of fibroblasts in the primary tumor and ascites. A) Pseudotime trajectory analysis showed that C0 and C1 were the origin of the cell lineage of fibroblasts, and C2 was at the end of the trajectory in the deep layer of the primary tumor (left). The expression of cancer-associated fibroblast (CAF) related genes reveals C0 (*CXCL12+),* C1 (*ACTA2+/S100A4+),* and C2 (*PDPN+/FN1+)* (middle). The msEMT genes were mainly yet differentially expressed in C0, C1, and C2 (right). B) In the ascites, eight sub-populations of fibroblasts were identified, and C1 and C3 were at the origin of the pseudotime trajectory, with C5 in the middle, and C6 and C7 at the end (left). The expression of CAF associated genes showed C1 (*VEGFA+),* C3 (*CXCL12+),* C5 (*ACTA2+/S100A4+*), C6 (*TNFRSF12A+*), and C7 (*S100A4+/FN1+)* (middle). The msEMT genes were mainly yet differentially expressed in C1, C3, C6, and C7, but were rarely expressed in C5, typical myofibroblastic CAF (right).

Comparing the gene expression profiles of fibroblasts in the primary tumor with those in ascites, we noted a dominance of up-regulated signals associated with the modulation of extracellular matrix within the primary tumor. Conversely, in ascites, there was significant up-regulation of signals related to heterotypic cell-cell adhesion and immune response (Supplementary Figure 5). These observations implicate an enhanced degree of interactions between fibroblasts and other cellular entities within the ascitic milieu compared to within the primary tumor. Therefore, we undertook a more comprehensive investigation to elucidate the dynamics of cell-to-cell interactions in the context of the primary tumor versus ascites.

### Interactions of fibroblasts in primary tumor and ascites

Upon conducting a cell-to-cell interaction analysis, we discerned that fibroblasts within primary tumors primarily orchestrate and respond to signals including Fibroblast Growth Factor (FGF), Midkine (MK), CXCL, and Pleiotrophin (PTN). This suggests a pivotal role for these fibroblasts in promoting tumor growth and progression in the primary tumor (Figure 6A). Conversely, fibroblasts within ascites predominantly receive signals such as the TNF-related weak inducer of apoptosis (TWEAK), and they primarily transmit signals encompassing Semaphorin 3 (SEMA3), CHEMERIN, MK, ANNEXIN, Complement, and Macrophage Migration Inhibitory Factor (MIF) (Figure 6B). These findings suggest that fibroblasts within ascites mainly foster migration and implantation of cancer cells by modulating the angiogenesis and immune evading within the peritoneal milieu, a new environment for cancer cells. It underscores that the observed changes in fibroblast signaling mirror the alterations in the dynamic interplay between cancer cells and fibroblasts throughout the metastatic cascade, underscoring their pivotal role in the peritoneal dissemination of GC.

**Figure 6.**
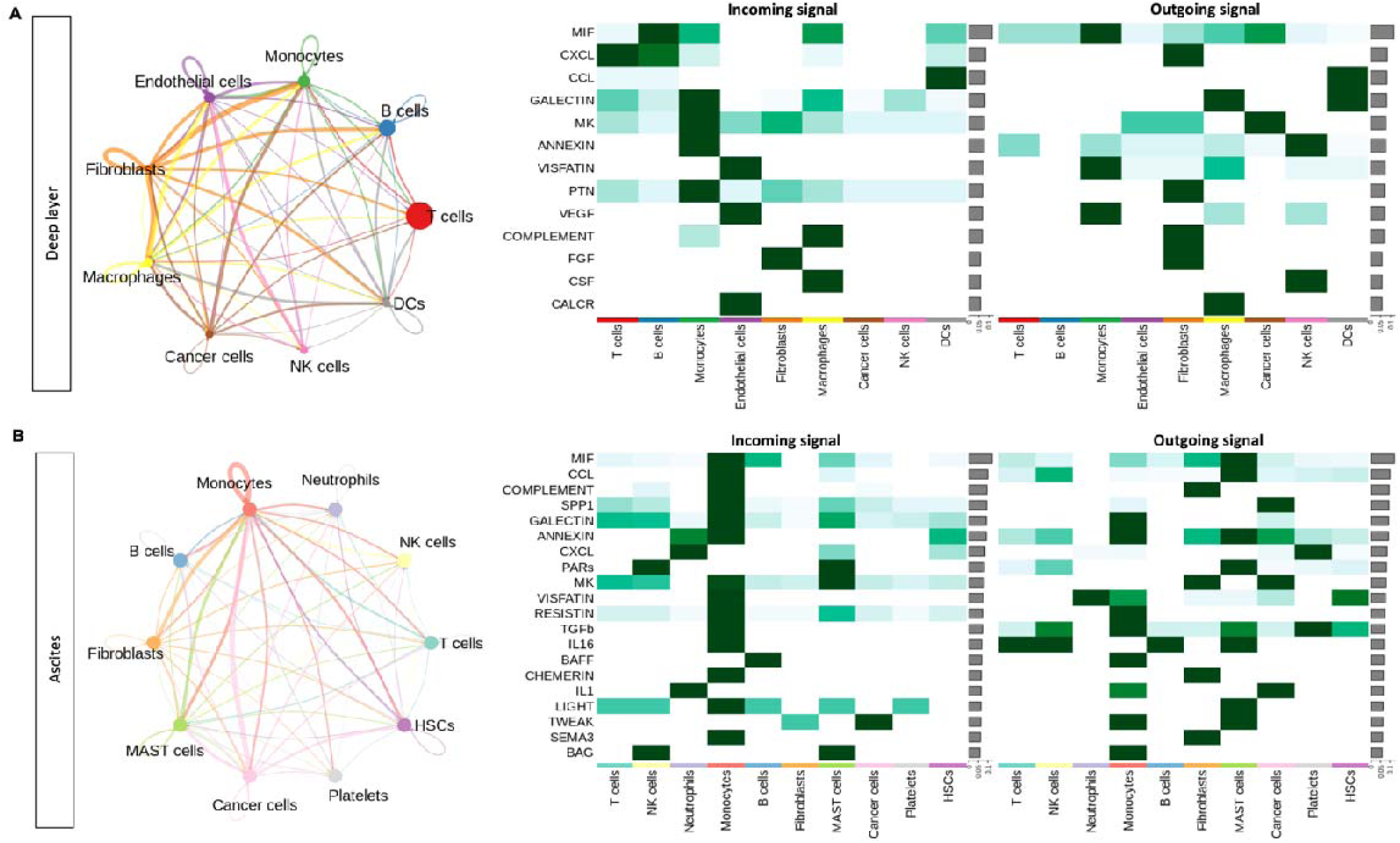
Interactions of fibroblasts with other cells in the primary tumor and ascites. A) In the deep layer of the primary tumors, the circle plot (left) showed the representation of T-cells as the most prominent, denoting their extensive presence across the cell groups. Concurrently, fibroblasts exhibited the thickest interconnections, underlining their heightened communication probability within the cellular milieu. Focusing on the fibroblasts, FGF, MK, CXCL, and PTN signals were mainly observed as incoming (middle) and outgoing (right) signals. B) In the ascites, a balanced cell population (similar circle sizes) was observed, and monocytes played a dominant role in intercellular communications (left). The TNF-related weak inducer of apoptosis (TWEAK) signal and Semaphorin 3 (SEMA3), CHEMERIN, MK, ANNEXIN, Complement, and Macrophage Migration Inhibitory Factor (MIF) were incoming (middle) and outgoing (right) signals in the fibroblasts in the ascites, respectively.

## Discussion

In this study, we identified metastatic GC specific genes whose expression patterns varied according to the metastatic routes: RTKs associated signals were up-regulated, while EMT associated signals were down-regulated in hematogenous metastasis. Conversely, EMT associated signals were up-regulated in peritoneal/ovarian metastases. These observations suggest that the molecular mechanisms driving GC metastasis may be route-specific, supporting previous report. [11] A recent large-scale genomic pan-cancer cohort study has provided invaluable insights into the overall differences between primary and metastatic solid cancers (early versus late stages of cancer), although the differences specific to metastatic routes were not thoroughly evaluated. [47] Our findings emphasize the necessity for future research and clinical strategies to not only consider the presence of metastasis (typically classified as M1 stage) but also the specific route of metastasis.

Additionally, we have identified a unique gene set, referred to as metastasis-specific epithelial-mesenchymal transition (msEMT) genes, which is specifically associated with metastatic GC tumors. This finding highlights a gap in previous research that primarily focused on primary tumors. [6, 8, 13] Interestingly, these genes were predominantly expressed in the tumor microenvironment, particularly in fibroblasts, rather than the cancer cells themselves. This suggests that GC metastasis is driven not just by genetic and epigenetic alterations within cancer cells but also by intricate interactions with the surrounding microenvironment. Cancer-associated fibroblasts (CAFs) play a crucial role in modulating cancer metastasis through extracellular matrix remodeling, growth factor production, and the regulation of angiogenesis and the immune system. [48, 49] It has been proposed that CAFs can exhibit distinct functional states, such as matrix-producing contractile phenotype (myo-CAFs), immuno- or vascular-modulating secretome (i-CAFs or v-CAFs). [48–51] In our study, msEMT genes were expressed in various fibroblast sub-populations, and the type of sub-populations predominantly expressing msEMT genes evolved during the progression of GC peritoneal metastasis. This finding suggests that msEMT genes contribute to the diversity and specificity of CAFs during GC peritoneal metastasis, promoting enhanced interactions with other cell types within ascites. Such dynamic interplay facilitates cancer cell migration and implantation by modulating angiogenesis and immune evasion, thereby highlighting the pivotal role for fibroblasts in the peritoneal dissemination of GC.

Another significant finding of our study is the association between the expression of metastatic tumor-specific genes in the primary tumor and clinical outcomes. A higher expression of msEMT genes in primary tumors was correlated with poor prognosis and demonstrated strong predictive performance for peritoneal/ovarian metastasis, exhibiting notable negative predictive value and specificity. This indicates that GC possesses metastatic potential even before the onset of peritoneal metastasis, and thus, msEMT genes could serve as potential biomarkers for risk assessment and for identifying GC patients who are less likely to develop peritoneal/ovarian metastasis. Furthermore, prior report have noted that the reprogramming of the tumor microenvironment plays a crucial role in primary tumor invasion of the stomach wall. [29] As the primary tumor invades the end of the stomach wall, cancer cells breach the serosa layer, leading to the spilling of cancer cells and microenvironment components together into the peritoneal space. This pre-arranged ecosystem offers a conducive environment for cancer cell dissemination and survival, providing a selective advantage in the peritoneal space. The immune landscape of the peritoneal space in GC patients exhibits distinctive characteristics, such as increased pro-angiogenic monocyte-like dendritic cells with reduced antigen-presenting capacity. [52] Given that the peritoneal cavity is a relatively sterile environment rarely exposed to invaders, [53] it may provide a less hostile milieu for GC, thus explaining why peritoneal metastasis is the most frequent metastatic or recurrent patterns in GC. Further single-cell level studies investigating the interactions between cancer cells and the tumor microenvironment, considering tumor location (e.g., primary tumor layers, free cells in ascites before peritoneal metastasis onset, and transplanted peritoneal metastatic tumors), may provide additional insights into the mechanisms of peritoneal metastasis.

## Conclusions

Our study has illuminated the route-specific mechanisms of GC metastasis and identified a unique gene set, the msEMT genes, associated specifically with metastatic GC. Furthermore, we underscored the pivotal role of fibroblasts within the tumor microenvironment, particularly those expressing msEMT genes, suggesting their contribution to the progression and specificity of peritoneal metastasis. These findings highlight the potential use of msEMT genes as biomarkers for risk assessment in the development of peritoneal/ovarian metastasis, and the need for further research to consider not just the presence but also the specific route of metastasis.

## List of abbreviations

GC: gastric cancer
TCGA: the cancer genome atlas
MSI: microsatellite instability
EBV: Epstein-Barr virus
CIN: chromosomal instability
GS: genomic stable
msEMT: metastasis-specific epithelial-mesenchymal transition
scRNA: single-cell RNA
FFPE: formalin-fixed paraffin-embedded
TNM: Tumor Node Metastasis
WTS: whole transcriptome sequencing
NMF: Non-negative Matrix Factorization
HR: hazard ratio
CI: confidence interval
EN: Elastic Net regression
GBM: Gradient Boosting Machine
RF: Random Forests
SVM: Support Vector Machine
PCANN: Probabilistic Classifier Artificial Neural Networks
DEA: Differential Expression Analysis
CAF: Cancer Associated Fibroblast
myo-CAF: myofibroblastic CAF
i-CAF: inflammatory CAF
v-CAF: vascular CAF

## Acknowledgement

The illustrations in figures were created with BioRender.com.

## Declarations

### Ethics approval and consent to participate

This study was approved by the Institutional Review Board (IRB) of Severance Hospital (4-2019-0188), Soonchunhyang University Bucheon Hospital (2022-09-008) and informed consent was received for all patients.

### Author contributions

Conceptualization: KTK, JEL, YYC

Methodology: KTK, YYC

Investigation: KTK, JEL, IC, JHC, YYC

Visualization: KTK, JEL, YYC

Funding acquisition: JEL, YYC

Project administration: KTK, JEL, IC, YYC

Supervision: KTK, YYC

Writing – original draft: KTK, JEL, YYC

Writing – review & editing: JHC, IC, KTK, JEL, YYC

### Data, Materials, and Code Availability

Raw data for the whole transcriptome sequencing (WTS) and single cell RNA-seq (scRNA-seq) reported in this manuscript have been deposited into a sequence read archive (SRA) database with BioProject accession number PRJNA973809 and PRJNA992126, respectively. The supported data which analyzed in this manuscript were publicly available from the GEO (www.ncbi.nlm.nih.gov/geo/) under accession numbers GSE167297 (scRNA-seq by layer of diffuse type of GC), GSE66229 (ACRG, primary GC) and GSE84437 (YCC, primary GC). Additionally, the code utilized for this analysis is not proprietary. All software and algorithms used in this study are freely accessible and are detailed in the Methods section. For more detailed information, please contact the corresponding author. (YYC)

### Consent for publication

Not applicable

---

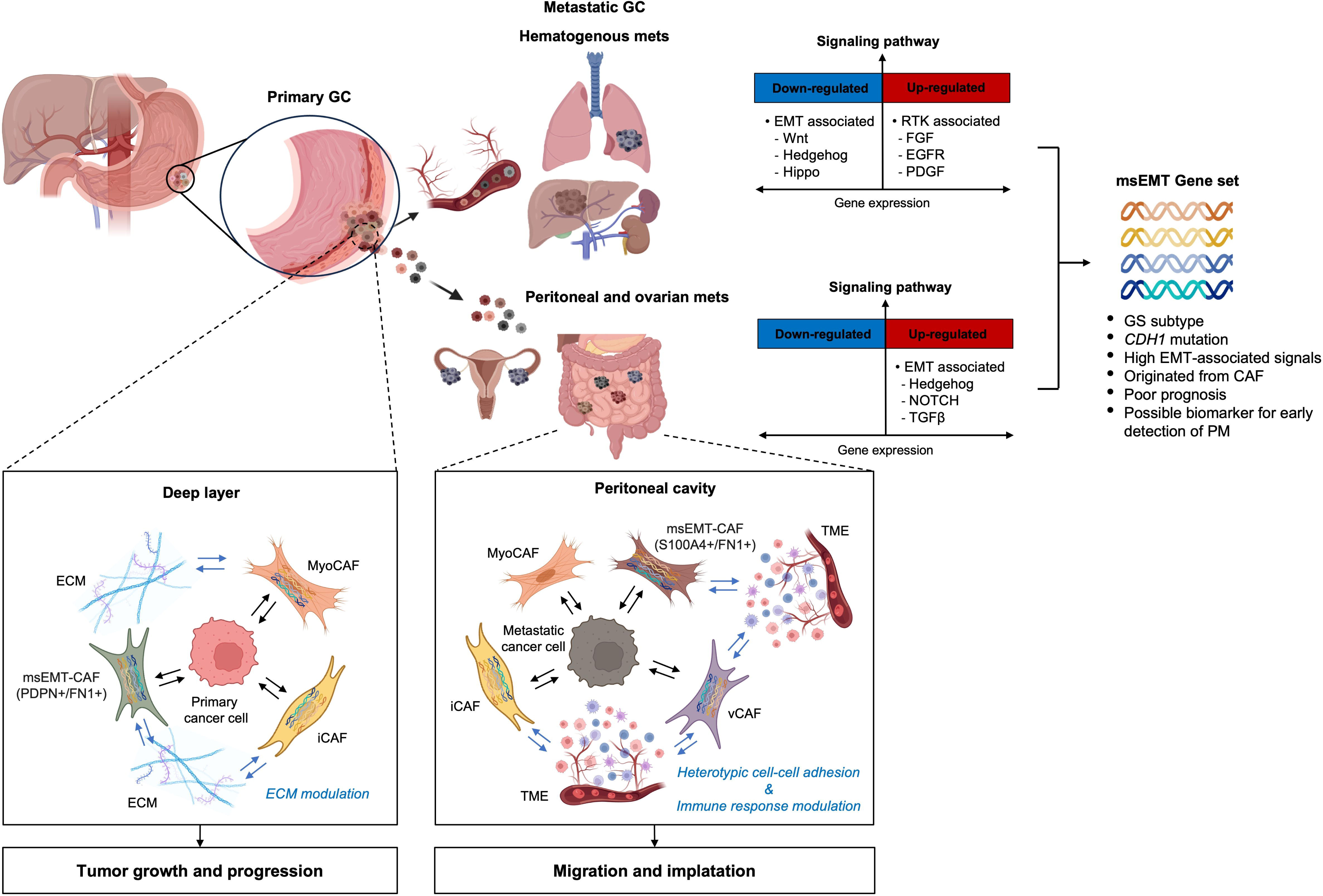

