## Supplementary Materials for "Multi-omics Identification and Route-Specific Characterization of Metastasis-specific EMT Genes and Their Microenvironmental Interactions"

**Supplementary Figure 1. Flow diagram for the analysis of the present study.**

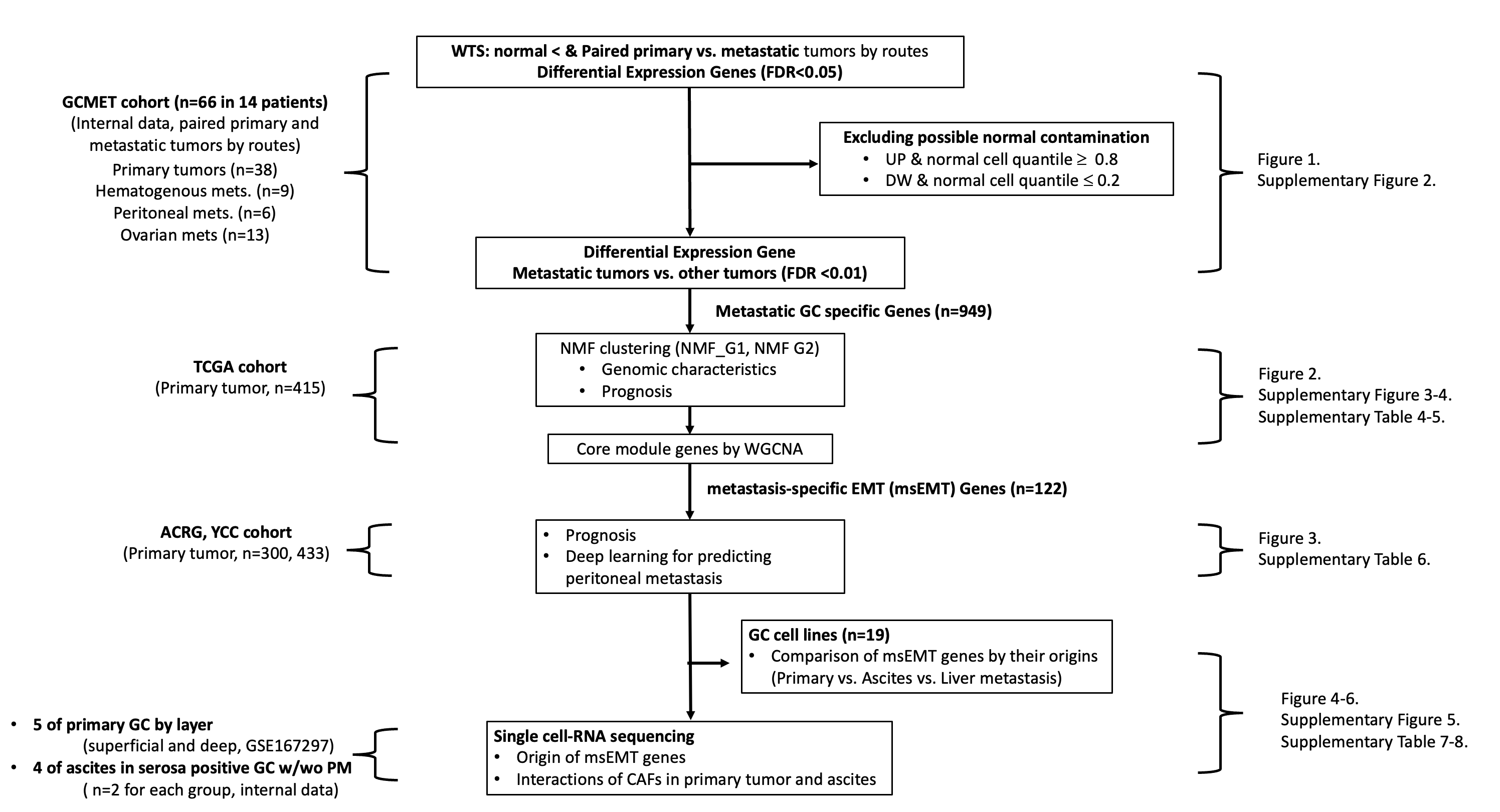

**Supplementary Figure 2. Gene set enrichment analysis for hedgehog pathway in hematogenous and peritoneal/ovarian metastasis specific genes.**

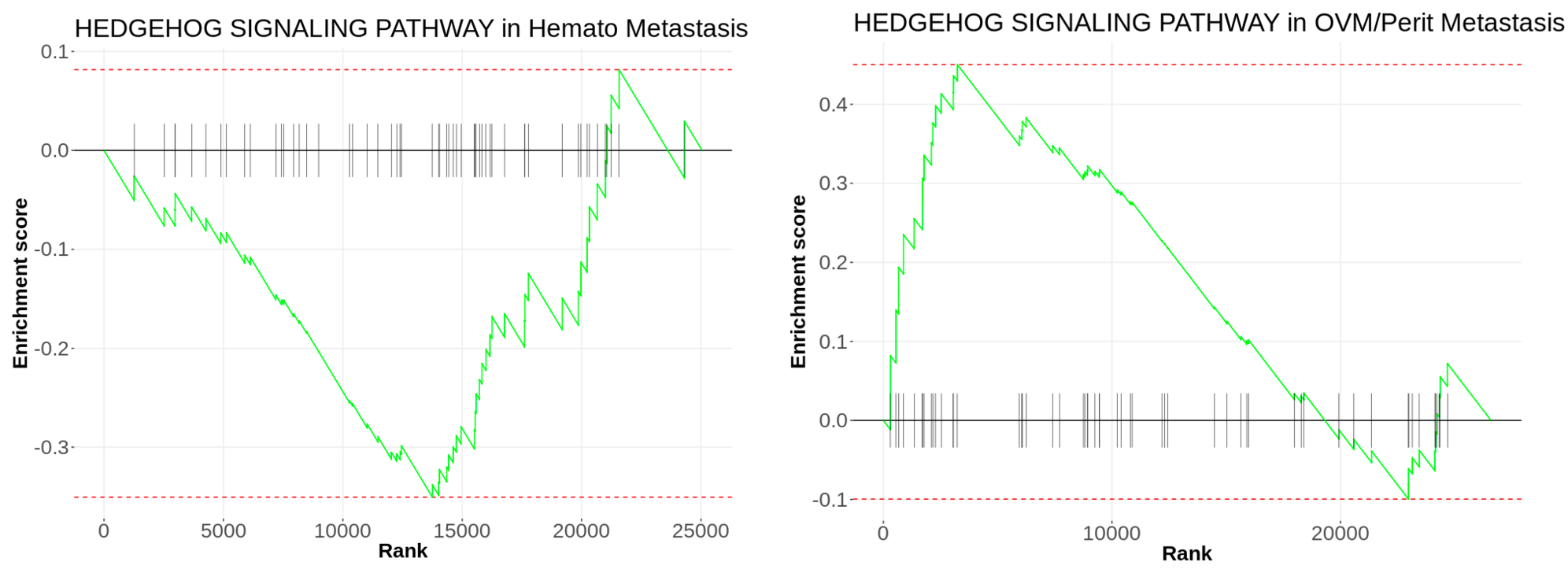

**Supplementary Figure 3.** **The characteristics of metastatic gastric cancer (GC) specific genes in The Cancer Genome Atlas (TCGA) cohort (n=415).** A) Non-negative matrix factorization (NMF) analysis, B) Comparing proportion of molecular subtypes of GC between NMF_G1 and NMF_G2, C) Overall survival by NMF-group in each molecular subtype of GC, D) oncoplot for somatic mutation by NMF-group.

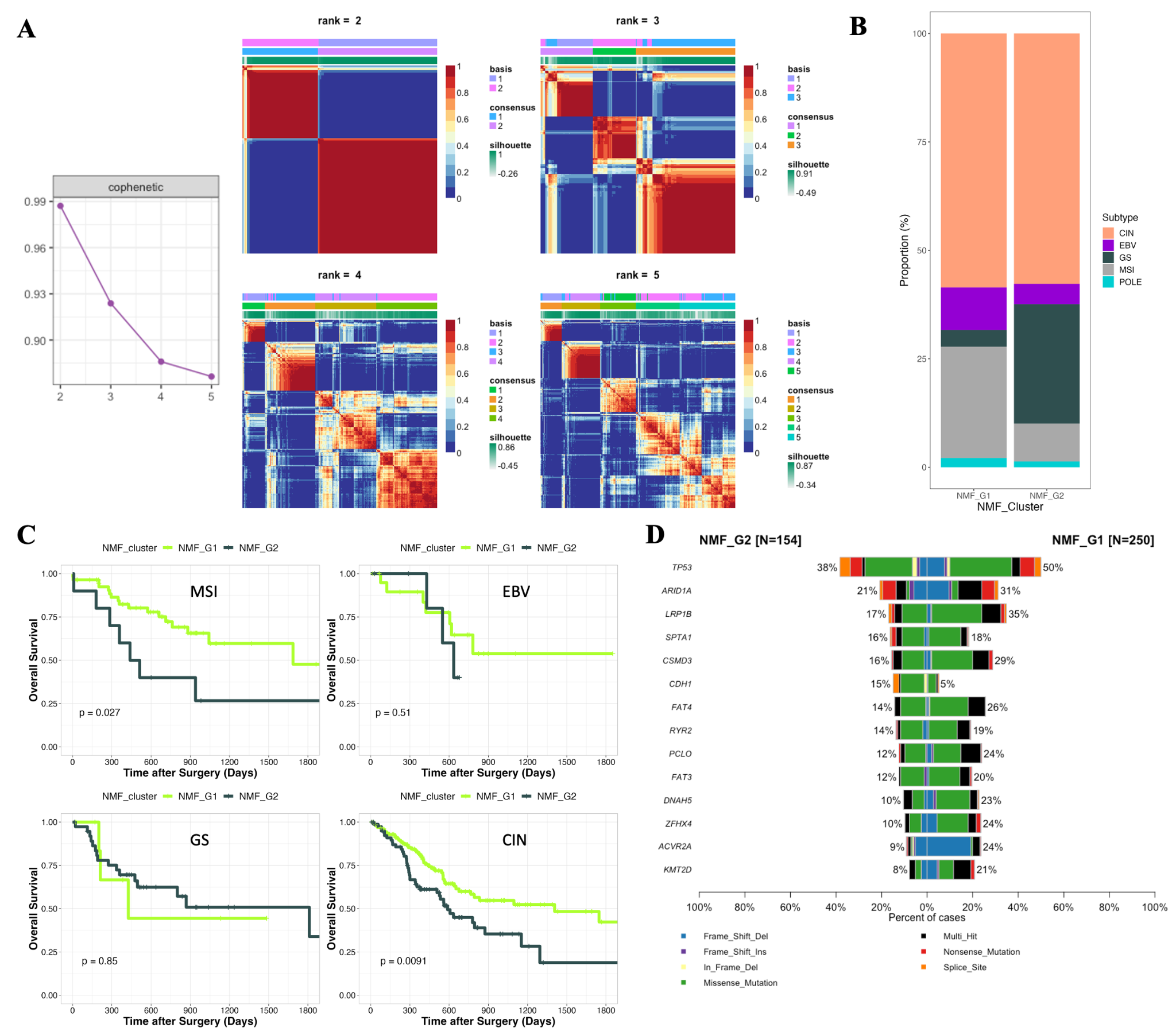

**Supplementary Figure 4. Weighted gene co-expression network analysis with metastatic gastric cancer specific genes in The Cancer Genome Atlas cohort.**

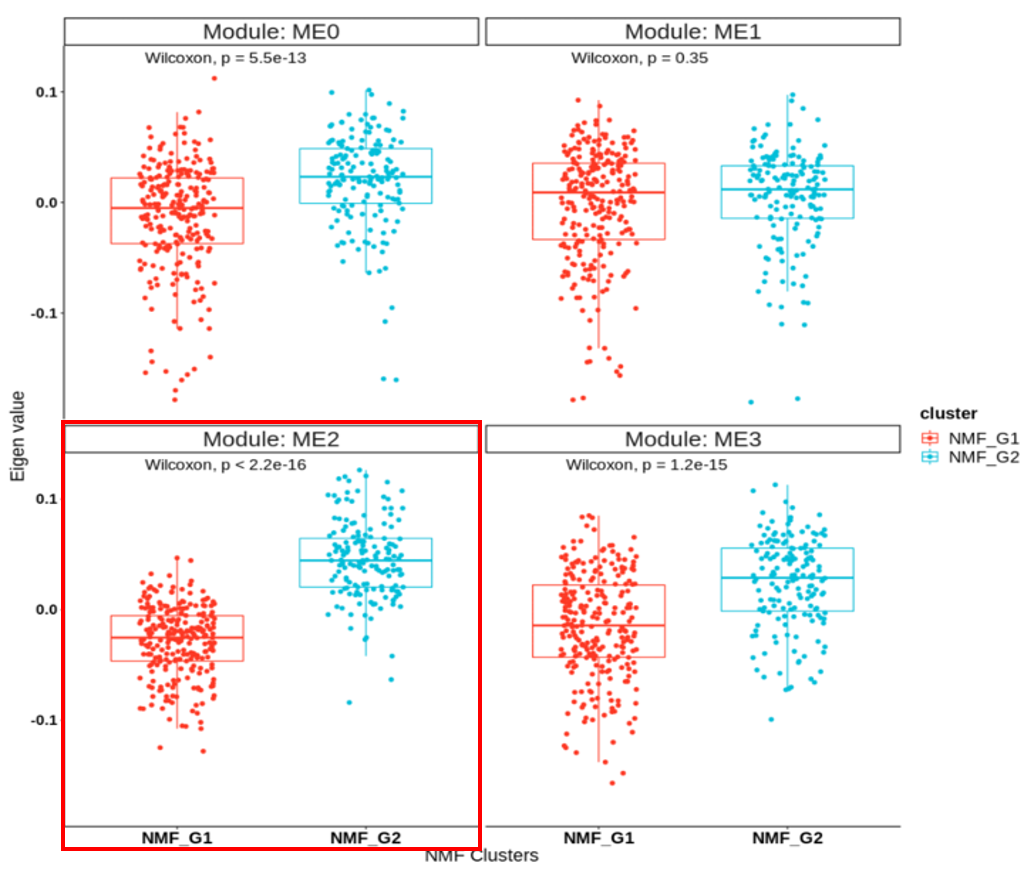

**Supplementary Figure 5. The over representation analysis by differential expression genes between fibroblasts within the primary tumor (deep layer) and ascites of patients with serosa positive gastric cancer.**

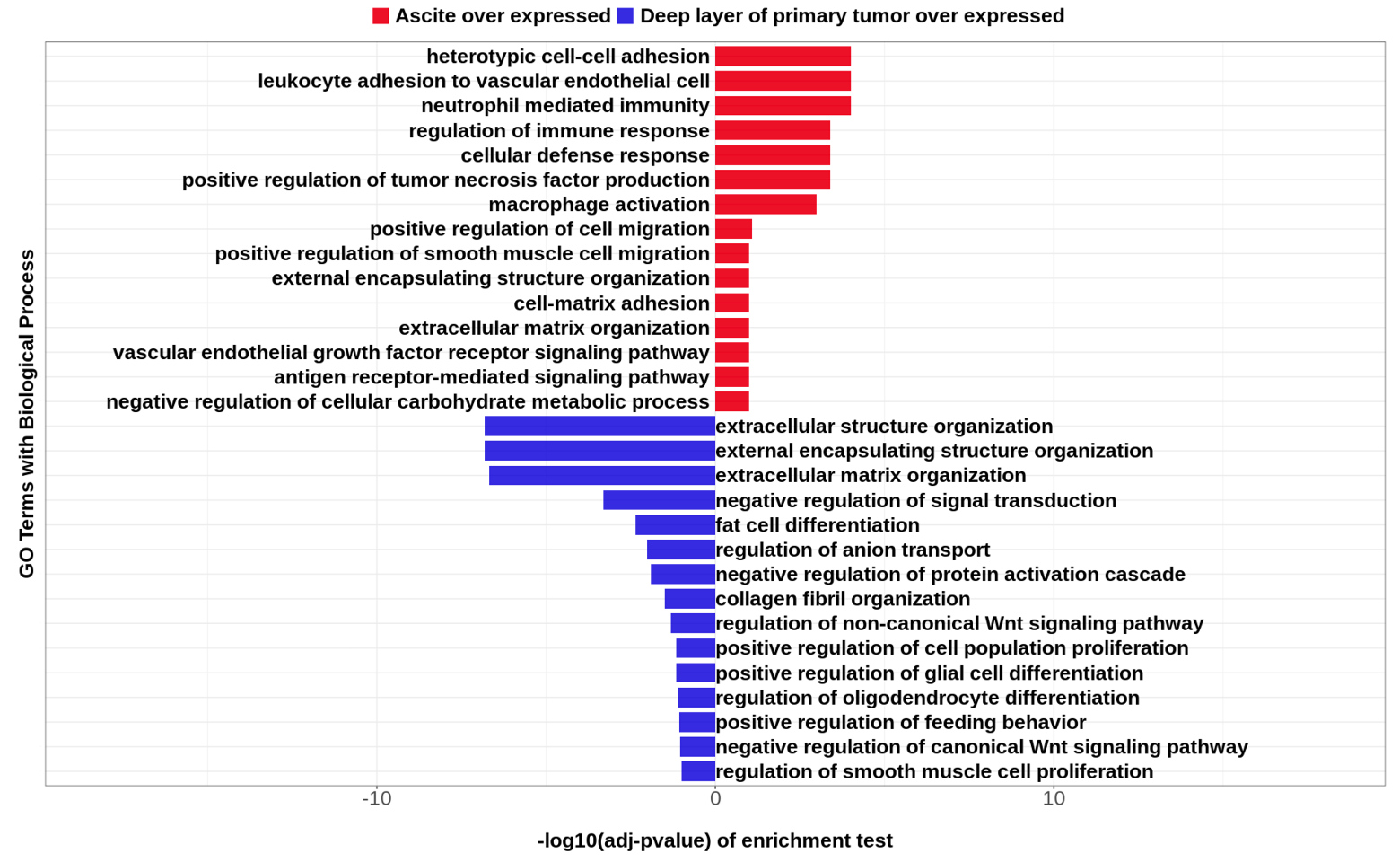

**Supplementary Table 1. Pathologic informations of enrolled patient with metastatic gastric cancer.**

| **PatientID** | **Age** | **Sex** | **TNM stage** | **Laeuren** | **Histology** | **MSI status** | **EBV status** |  |
| --- | --- | --- | --- | --- | --- | --- | --- | --- |
| GCM01 | 68 | M | IA | Intestinal | MD | MSS | negative |  |
| GCM02 | 73 | M | IIB | Intestinal | MD | MSS | negative |  |
| GCM03 | 55 | M | IIB | Intestinal | MD | MSS | negative |  |
| GCM04 | 43 | M | IIIB | Intestinal | MD | MSS | negative |  |
| GCM05 | 47 | F | IIIA | Mixed | PD with mucin pool | MSS | negative |  |
| GCM06 | 29 | F | IIIB | Intestinal | PD | MSS | negative |  |
| GCM07 | 43 | F | IIIB | Diffuse | Mucinous | MSS | negative |  |
| GCM08 | 38 | F | IIIC | Diffuse | SRC | MSS | negative |  |
| GCM09 | 47 | F | IIIC | Diffuse | SRC with mucin pool | MSS | negative |  |
| GCM10 | 33 | M | IIIC | Diffuse | PD | MSS | negative |  |
| GCM12 | 53 | F | IIIA | Diffuse | SRC | MSS | negative |  |
| GCM13 | 38 | F | IV | Diffuse | PD with SRC | MSS | negative |  |
| GCM14 | 56 | F | IIIB | Diffuse | SRC | MSS | negative |  |
| GCM15 | 72 | M | IIIB | Intestinal | MD | MSS | negative |  |
| **PatientID** | **Age** | **Sex** | **TNM stage** | **Laeuren** | **Histology** | **Metastasis** | **MSI status** | **EBV status** |
| GCMsc_01 | 71 | F | IIIC | Diffuse | SRC | CY- | MSS | negative |
| GCMsc_02 | 51 | M | IV | Diffuse | SRC | CY+, PM+ | MSS | negative |
| GCMsc_03 | 64 | M | IV | Diffuse | PD | CY+, PM+ | MSS | negative |
| GCMsc_04 | 63 | M | IIIA | Diffuse | PD | CY- | MSS | negative |

MD; moderate differentiated, PD; poorly differentiated, SRC; signet ring cell carcinoma, CY: cytology, PM: peritoneal metastasis, MSS: microsatellite stable, EBV; Epstein Barr Virus

**Supplementary Table 2. Information for tumor samples in each patient with metastatic gastric cancer.**

| **PatientID** | **SampleID** | **Met Type** | **Met timing** | **Exposure to chemotherapy** | **Lauren classification** | **Purity*** | **Microsatellite instability status#** |
| --- | --- | --- | --- | --- | --- | --- | --- |
| GCM01 | GCM01_T | primary | synchronous | naïve | Intestinal | 0.5 | microsatellite stable |
| GCM01 | GCM01_liver | hematogenous | metachronous | naïve | Intestinal | 0.8 | microsatellite stable |
| GCM02 | GCM02_T1 | primary | synchronous | naïve | Intestinal | 0.25 | microsatellite stable |
| GCM02 | GCM02_T2 | primary | synchronous | naïve | Intestinal | 0.3 | microsatellite stable |
| GCM02 | GCM02_liver1 | hematogenous | metachronous | exposed | Intestinal | 0.7 | microsatellite stable |
| GCM02 | GCM02_liver2 | hematogenous | metachronous | exposed | Intestinal | 0.7 | microsatellite stable |
| GCM02 | GCM02_liver3 | hematogenous | metachronous | exposed | Intestinal | 0.6 | microsatellite stable |
| GCM03 | GCM03_T | primary | synchronous | naïve | Intestinal | 0.3 | microsatellite stable |
| GCM03 | GCM03_liver1 | hematogenous | metachronous | exposed | Intestinal | 0.7 | microsatellite stable |
| GCM03 | GCM03_liver2 | hematogenous | metachronous | exposed | Intestinal | 0.6 | microsatellite stable |
| GCM04 | GCM04_T1 | primary | synchronous | naïve | Intestinal | 0.45 | microsatellite stable |
| GCM04 | GCM04_T2 | primary | synchronous | naïve | Intestinal | 0.2 | microsatellite stable |
| GCM04 | GCM04_T3 | primary | synchronous | naïve | Intestinal | 0.3 | microsatellite stable |
| GCM04 | GCM04_lung | hematogenous | metachronous | exposed | Intestinal | 0.8 | microsatellite stable |
| GCM05 | GCM05_T1 | primary | synchronous | naïve | Mixed | 0.15 | microsatellite stable |
| GCM05 | GCM05_T2 | primary | synchronous | naïve | Mixed | 0.1 | microsatellite stable |
| GCM05 | GCM05_T3 | primary | synchronous | naïve | Mixed | 0.1 | microsatellite stable |
| GCM05 | GCM05_perit1 | peritoneum | metachronous | exposed | Mixed | 0.05 | microsatellite stable |
| GCM05 | GCM05_perit2 | peritoneum | metachronous | exposed | Mixed | 0.05 | microsatellite stable |
| GCM06 | GCM06_T1a | primary | synchronous | naïve | Intestinal | 0.3 | microsatellite stable |
| GCM06 | GCM06_T1b | primary | synchronous | naïve | Intestinal | 0.1 | microsatellite stable |
| GCM06 | GCM06_T2 | primary | synchronous | naïve | Intestinal | 0.05 | microsatellite stable |
| GCM06 | GCM06_T3 | primary | synchronous | naïve | Intestinal | 0.05 | microsatellite stable |
| GCM06 | GCM06_ovary1 | ovary | metachronous | exposed | Intestinal | 0.15 | microsatellite stable |
| GCM06 | GCM06_ovary2 | ovary | metachronous | exposed | Intestinal | 0.3 | microsatellite stable |
| GCM07 | GCM07_T1a | primary | synchronous | naïve | Diffuse | 0.1 | microsatellite stable |
| GCM07 | GCM07_T1b | primary | synchronous | naïve | Diffuse | 0.6 | microsatellite stable |
| GCM07 | GCM07_T2 | primary | synchronous | naïve | Diffuse | 0.6 | microsatellite stable |
| GCM07 | GCM07_T3 | primary | synchronous | naïve | Diffuse | 0.2 | microsatellite stable |
| GCM07 | GCM07_ovary | ovary | metachronous | exposed | Diffuse | 0.6 | microsatellite stable |
| GCM08 | GCM08_T1 | primary | synchronous | naïve | Diffuse | 0.9 | microsatellite stable |
| GCM08 | GCM08_T2 | primary | synchronous | naïve | Diffuse | 0.5 | microsatellite stable |
| GCM08 | GCM08_T3 | primary | synchronous | naïve | Diffuse | 0.3 | microsatellite stable |
| GCM08 | GCM08_ovary1 | ovary | metachronous | exposed | Diffuse | 0.15 | microsatellite stable |
| GCM08 | GCM08_ovary2 | ovary | metachronous | exposed | Diffuse | 0.15 | microsatellite stable |
| GCM09 | GCM09_T1 | primary | synchronous | naïve | Diffuse | 0.8 | microsatellite stable |
| GCM09 | GCM09_T2 | primary | synchronous | naïve | Diffuse | 0.5 | microsatellite stable |
| GCM09 | GCM09_T3 | primary | synchronous | naïve | Diffuse | 0.4 | microsatellite stable |
| GCM09 | GCM09_perit | peritoneum | metachronous | exposed | Diffuse | 0.05 | microsatellite stable |
| GCM09 | GCM09_ovary1 | ovary | metachronous | exposed | Diffuse | 0.4 | microsatellite stable |
| GCM09 | GCM09_ovary2 | ovary | metachronous | exposed | Diffuse | 0.3 | microsatellite stable |
| GCM10 | GCM10_T1a | primary | synchronous | exposed | Diffuse | 0.3 | microsatellite stable |
| GCM10 | GCM10_T1b | primary | synchronous | exposed | Diffuse | 0.05 | microsatellite stable |
| GCM10 | GCM10_T2 | primary | synchronous | exposed | Diffuse | 0.01 | microsatellite stable |
| GCM10 | GCM10_T3 | primary | synchronous | exposed | Diffuse | 0.05 | microsatellite stable |
| GCM10 | GCM10_liver | hematogenous | synchronous | exposed | Diffuse | 0.4 | microsatellite stable |
| GCM12 | GCM12_T1a | primary | synchronous | naïve | Diffuse | 0.1 | microsatellite stable |
| GCM12 | GCM12_T1b | primary | synchronous | naïve | Diffuse | 0.1 | microsatellite stable |
| GCM12 | GCM12_T2 | primary | synchronous | naïve | Diffuse | 0.05 | microsatellite stable |
| GCM12 | GCM12_perit | peritoneum | synchronous | naïve | Diffuse | 0.02 | microsatellite stable |
| GCM12 | GCM12_ovary1 | ovary | synchronous | naïve | Diffuse | 0.3 | microsatellite stable |
| GCM12 | GCM12_ovary2 | ovary | synchronous | naïve | Diffuse | 0.2 | microsatellite stable |
| GCM13 | GCM13_T1 | primary | synchronous | naïve | Diffuse | 0.4 | microsatellite stable |
| GCM13 | GCM13_T2 | primary | synchronous | naïve | Diffuse | 0.2 | microsatellite stable |
| GCM13 | GCM13_perit | peritoneum | synchronous | naïve | Diffuse | 0.3 | microsatellite stable |
| GCM13 | GCM13_ovary1 | ovary | synchronous | naïve | Diffuse | 0.05 | microsatellite stable |
| GCM13 | GCM13_ovary2 | ovary | synchronous | naïve | Diffuse | 0.3 | microsatellite stable |
| GCM14 | GCM14_T1 | primary | synchronous | naïve | Diffuse | 0.27 | microsatellite stable |
| GCM14 | GCM14_T2 | primary | synchronous | naïve | Diffuse | 0.02 | microsatellite stable |
| GCM14 | GCM14_perit | peritoneum | metachronous | exposed | Diffuse | 0.9 | microsatellite stable |
| GCM14 | GCM14_ovary1 | ovary | metachronous | exposed | Diffuse | 0.15 | microsatellite stable |
| GCM14 | GCM14_ovary2 | ovary | metachronous | exposed | Diffuse | 0.2 | microsatellite stable |
| GCM15 | GCM15_T1 | primary | synchronous | naïve | Intestinal | 0.32 | microsatellite stable |
| GCM15 | GCM15_T2 | primary | synchronous | naïve | Intestinal | 0.25 | microsatellite stable |
| GCM15 | GCM15_T3 | primary | synchronous | naïve | Intestinal | 0.1 | microsatellite stable |
| GCM15 | GCM15_liver | hematogenous | metachronous | exposed | Intestinal | 0.3 | microsatellite stable |

* estimated by ASCAT

\# estimated by MSI Sensor

**Supplementary Table 3. Information for scRNA-seq.**

| **Sample** | **Estimated Number of Cells before filtering** | **Estimated Number of Cells after QC and artifical doublet filtering** | **Mean Reads per Cell** | **Median Genes per Cell** | **Median UMI Counts per Cell** | **Reads Mapped to Genome** | **Valid barcode** | **Q30 Bases in UMI** | **Sequencing platform** |
| --- | --- | --- | --- | --- | --- | --- | --- | --- | --- |
| GCMsc_01 | 12,693 | 9,981 | 32,324 | 1,745 | 4,748 | 94.70% | 97.60% | 97.80% | Illumina Hiseq |
| GCMsc_02 | 10,145 | 8,514 | 39,989 | 1,628 | 4,541 | 96.70% | 97.80% | 97.70% | Illumina Hiseq |
| GCMsc_03 | 9,288 | 7,791 | 46,444 | 1,650 | 4,314 | 97.30% | 98.10% | 97.90% | Illumina Hiseq |
| GCMsc_04 | 13,233 | 7,091 | 32,607 | 1,741 | 5,001 | 96.90% | 97.70% | 97.50% | Illumina Hiseq |

**Supplementary Table 4. The 122 of metastasis specific Epthelial-Mesenchymal Transition (msEMT) genes and the associations to well-established gene-sets.**

| **Symbol** | **DEG_type** | **Immune** | **EMT_up** | **EMT_dw** | **MSI_UP** | **Proliferation** | **Gastric_tissue** | **Cancer_driver_genes** |
| --- | --- | --- | --- | --- | --- | --- | --- | --- |
| ABCC9 | Hemato_dw | No | No | No | No | No | No | No |
| ADAMTSL3 | Hemato_dw | No | No | No | No | No | No | No |
| ADPRH | Hemato_dw | No | No | No | No | No | No | No |
| AKAP6 | Hemato_dw | No | No | No | No | No | No | No |
| ANKRD53 | Hemato_dw | No | No | No | No | No | No | No |
| ARHGAP20 | Hemato_dw | No | No | No | No | No | No | No |
| ARHGAP22 | Hemato_dw | No | No | No | No | No | No | No |
| ATP1B2 | Hemato_dw | No | No | No | No | No | No | No |
| BCHE | Hemato_dw | No | No | No | No | No | No | No |
| C1QTNF7 | Hemato_dw | No | No | No | No | No | No | No |
| CACNA2D1 | Hemato_dw | No | No | No | No | No | No | No |
| CASQ1 | Hemato_dw | No | No | No | No | No | No | No |
| CCDC136 | Hemato_dw | No | No | No | No | No | No | No |
| CDON | Hemato_dw | No | No | No | No | No | No | No |
| CMYA5 | Hemato_dw | No | No | No | No | No | No | No |
| CNKSR2 | Hemato_dw | No | No | No | No | No | No | No |
| CORO2B | Hemato_dw | No | No | No | No | No | No | No |
| DIXDC1 | Hemato_dw | No | No | No | No | No | No | No |
| DLG2 | Hemato_dw | No | No | No | No | No | No | No |
| DLG4 | Hemato_dw | No | No | No | No | No | No | No |
| DOK6 | Hemato_dw | No | No | No | No | No | No | No |
| DSEL | Hemato_dw | No | No | No | No | No | No | No |
| DYNC1I1 | Hemato_dw | No | No | No | No | No | No | No |
| EBF3 | Hemato_dw | No | No | No | No | No | No | No |
| EDA2R | Hemato_dw | No | No | No | No | No | No | No |
| EPM2A | Hemato_dw | No | No | No | No | No | No | No |
| ERG | Hemato_dw | No | No | No | No | No | No | No |
| FAM124A | Hemato_dw | No | No | No | No | No | No | No |
| FAM13C | Hemato_dw | No | No | No | No | No | No | No |
| FBXL7 | Hemato_dw | No | No | No | No | No | No | No |
| GLI1 | Hemato_dw | No | No | No | No | No | No | No |
| GLI3 | Hemato_dw | No | No | No | No | No | No | No |
| GPM6A | Hemato_dw | No | No | No | No | No | No | No |
| GREB1 | Hemato_dw | No | No | No | No | No | No | No |
| HHIPL1 | Hemato_dw | No | No | No | No | No | No | No |
| HOXA4 | Hemato_dw | No | No | No | No | No | No | No |
| HOXA5 | Hemato_dw | No | No | No | No | No | No | No |
| KATNAL1 | Hemato_dw | No | No | No | No | No | No | No |
| KCNMA1 | Hemato_dw | No | No | No | No | No | No | No |
| KIAA1614 | Hemato_dw | No | No | No | No | No | No | No |
| KIAA1755 | Hemato_dw | No | No | No | No | No | No | No |
| LRRC4C | Hemato_dw | No | No | No | No | No | No | No |
| LRRN2 | Hemato_dw | No | No | No | No | No | No | No |
| MAP1A | Hemato_dw | No | No | No | No | No | No | No |
| MAP7D3 | Hemato_dw | No | No | No | No | No | No | No |
| MAP9 | Hemato_dw | No | No | No | No | No | No | No |
| MDGA1 | Hemato_dw | No | No | No | No | No | No | No |
| MKX | Hemato_dw | No | No | No | No | No | No | No |
| MN1 | Hemato_dw | No | No | No | No | No | No | No |
| MPPED2 | Hemato_dw | No | No | No | No | No | No | No |
| MYCT1 | Hemato_dw | No | No | No | No | No | No | No |
| NACAD | Hemato_dw | No | No | No | No | No | No | No |
| NFATC4 | Hemato_dw | No | No | No | No | No | No | No |
| NOVA2 | Hemato_dw | No | No | No | No | No | No | No |
| NRXN2 | Hemato_dw | No | No | No | No | No | No | No |
| NUDT10 | Hemato_dw | No | No | No | No | No | No | No |
| PCDH9 | Hemato_dw | No | No | No | No | No | No | No |
| PCDHB4 | Hemato_dw | No | No | No | No | No | No | No |
| PCDHB5 | Hemato_dw | No | No | No | No | No | No | No |
| PCDHGA3 | Hemato_dw | No | No | No | No | No | No | No |
| PDE8B | Hemato_dw | No | No | No | No | No | No | No |
| PHACTR1 | Hemato_dw | No | No | No | No | No | No | No |
| PLP1 | Hemato_dw | No | No | No | No | No | No | No |
| PLXNA4 | Hemato_dw | No | No | No | No | No | No | No |
| PREX2 | Hemato_dw | No | No | No | No | No | No | No |
| PRICKLE1 | Hemato_dw | No | No | No | No | No | No | No |
| PRRX1 | Hemato_dw | No | No | No | No | No | No | No |
| PYGM | Hemato_dw | No | No | No | No | No | No | No |
| RASGRF2 | Hemato_dw | No | No | No | No | No | No | No |
| RNF165 | Hemato_dw | No | No | No | No | No | No | No |
| RUNX1T1 | Hemato_dw | No | No | No | No | No | No | No |
| SMAD9 | Hemato_dw | No | No | No | No | No | No | No |
| SMARCD3 | Hemato_dw | No | No | No | No | No | No | No |
| SORCS2 | Hemato_dw | No | No | No | No | No | No | No |
| SSC5D | Hemato_dw | No | No | No | No | No | No | No |
| TDRD10 | Hemato_dw | No | No | No | No | No | No | No |
| TMEM132E | Hemato_dw | No | No | No | No | No | No | No |
| TNN | Hemato_dw | No | No | No | No | No | No | No |
| TRIM61 | Hemato_dw | No | No | No | No | No | No | No |
| TSHZ3 | Hemato_dw | No | No | No | No | No | No | No |
| TTLL7 | Hemato_dw | No | Yes | No | No | No | No | No |
| UNC5C | Hemato_dw | No | No | No | No | No | No | No |
| USP51 | Hemato_dw | No | No | No | No | No | No | No |
| VGLL3 | Hemato_dw | No | No | No | No | No | No | No |
| WNK3 | Hemato_dw | No | No | No | No | No | No | No |
| WNT9B | Hemato_dw | No | No | No | No | No | No | No |
| ZFHX4 | Hemato_dw | No | No | No | No | No | No | No |
| ZFP28 | Hemato_dw | No | No | No | No | No | No | No |
| ZFPM2 | Hemato_dw | No | Yes | No | No | No | No | No |
| ZNF135 | Hemato_dw | No | No | No | No | No | No | No |
| ZNF471 | Hemato_dw | No | No | No | No | No | No | No |
| ZNF483 | Hemato_dw | No | No | No | No | No | No | No |
| ZNF521 | Hemato_dw | No | No | No | No | No | No | No |
| ZNF660 | Hemato_dw | No | No | No | No | No | No | No |
| ANKRD6 | ovm_up | No | No | No | No | No | No | No |
| C3orf70 | ovm_up | No | No | No | No | No | No | No |
| CAND2 | Hemato_dw/ovm_up | No | No | No | No | No | No | No |
| CPT1C | Hemato_dw/ovm_up | No | No | No | No | No | No | No |
| FAM110B | ovm_up | No | No | No | No | No | No | No |
| FGD5 | ovm_up | No | No | No | No | No | No | No |
| FHOD3 | Hemato_dw/ovm_up | No | No | No | No | No | No | No |
| FMO1 | ovm_up | No | No | No | No | No | No | No |
| GPR135 | Hemato_dw/ovm_up | No | No | No | No | No | No | No |
| IGDCC4 | ovm_up | No | No | No | No | No | No | No |
| KIRREL3 | ovm_up | No | No | No | No | No | No | No |
| MAGEL2 | ovm_up | No | No | No | No | No | No | No |
| MAGI2 | Hemato_dw/ovm_up | No | No | No | No | No | No | No |
| MYOCD | ovm_up | No | No | No | No | No | No | No |
| MYOZ3 | ovm_up | No | No | No | No | No | No | No |
| PDE1B | ovm_up | No | No | No | No | No | No | No |
| PKNOX2 | ovm_up | No | No | No | No | No | No | No |
| PPFIA2 | ovm_up | No | No | No | No | No | No | No |
| PRKD1 | Hemato_dw/ovm_up | No | Yes | No | No | No | No | No |
| SHROOM4 | ovm_up | No | No | No | No | No | No | No |
| SYT15 | ovm_up | No | No | No | No | No | No | No |
| TBC1D9 | ovm_up | No | No | No | No | No | No | No |
| TTC7B | ovm_up | No | No | No | No | No | No | No |
| YPEL4 | Hemato_dw/ovm_up | No | No | No | No | No | No | No |
| FAM124B | perit_up | No | No | No | No | No | No | No |
| NLGN4X | Hemato_dw/perit_up | No | No | No | No | No | No | No |
| GFRA2 | Hemato_dw/perit_up | No | No | No | No | No | No | No |
| CCDC89 | perit_up | No | No | No | No | No | No | No |

**Supplementary Table 5. The comparison of metastasis specific Epithelial-Mesenchymal Transition (msEMT) genes' expression between peritoneal or ovarian metastasis and primary tumors.**

| Gene | Primary_Mean | Perit_OVM_Mean | P_value | log2FC |
| --- | --- | --- | --- | --- |
| ABCC9 | 12.15799085 | 12.0638836 | 0.64835204 | -0.0941073 |
| ADAMTSL3 | 10.97001729 | 11.37766959 | 0.03573518 | 0.4076523 |
| ADPRH | 9.360100686 | 9.414778034 | 0.8083513 | 0.05467735 |
| AKAP6 | 11.34322466 | 11.34173585 | 0.995666063 | -0.0014888 |
| ANKRD53 | 8.136873868 | 8.60439967 | 0.071093102 | 0.4675258 |
| ANKRD6 | 9.920055949 | 10.63897444 | 0.002509014 | 0.71891849 |
| ARHGAP20 | 9.248097722 | 9.747769747 | 0.071929627 | 0.49967203 |
| ARHGAP22 | 9.160959076 | 10.1712667 | 4.92E-07 | 1.01030763 |
| ATP1B2 | 9.733985743 | 10.60865788 | 0.004249332 | 0.87467214 |
| BCHE | 9.128133162 | 9.191914486 | 0.873718207 | 0.06378132 |
| C1QTNF7 | 10.32757204 | 10.56187473 | 0.153535214 | 0.2343027 |
| C3orf70 | 10.42579712 | 11.05548256 | 0.00100851 | 0.62968544 |
| CACNA2D1 | 12.00667347 | 12.48930268 | 0.012182465 | 0.48262922 |
| CAND2 | 9.700002267 | 10.50950237 | 0.003954124 | 0.80950011 |
| CASQ1 | 5.84000461 | 6.13224083 | 0.556788427 | 0.29223622 |
| CCDC136 | 8.877483847 | 9.622717254 | 0.016129334 | 0.74523341 |
| CCDC89 | 8.676173713 | 9.183153493 | 0.004118775 | 0.50697978 |
| CDON | 10.19772343 | 11.23673649 | 0.006913054 | 1.03901306 |
| CMYA5 | 11.0463048 | 11.54088688 | 0.032263478 | 0.49458208 |
| CNKSR2 | 10.80440789 | 11.17433019 | 0.398152668 | 0.3699223 |
| CORO2B | 8.875884982 | 9.937251038 | 8.31E-05 | 1.06136606 |
| CPT1C | 9.444341989 | 10.67038262 | 8.64E-06 | 1.22604063 |
| DIXDC1 | 8.873458751 | 9.244110956 | 0.051164374 | 0.3706522 |
| DLG2 | 10.45397974 | 11.03342142 | 0.011513851 | 0.57944168 |
| DLG4 | 9.768822635 | 10.86957653 | 1.94E-06 | 1.10075389 |
| DOK6 | 11.0417987 | 11.37130779 | 0.032517301 | 0.32950909 |
| DSEL | 10.76028846 | 11.26271815 | 0.00458019 | 0.50242969 |
| DYNC1I1 | 9.241702902 | 9.660894227 | 0.185217843 | 0.41919132 |
| EBF3 | 10.01920546 | 10.60330303 | 0.058369751 | 0.58409757 |
| EDA2R | 8.553708674 | 8.757802393 | 0.3935199 | 0.20409372 |
| EPM2A | 9.935467127 | 10.1634548 | 0.217450402 | 0.22798767 |
| ERG | 11.05440166 | 11.38479086 | 0.077014279 | 0.3303892 |
| FAM110B | 9.008902309 | 9.983735089 | 0.000303784 | 0.97483278 |
| FAM124A | 9.421442783 | 9.530686827 | 0.681428524 | 0.10924404 |
| FAM124B | 8.440301257 | 8.586161936 | 0.676592192 | 0.14586068 |
| FAM13C | 8.645933159 | 9.199526421 | 0.003686938 | 0.55359326 |
| FBXL7 | 10.10338925 | 11.2721002 | 0.000280817 | 1.16871095 |
| FGD5 | 11.12717827 | 11.69418249 | 0.003893772 | 0.56700422 |
| FHOD3 | 9.23757003 | 10.1487544 | 0.000216901 | 0.91118437 |
| FMO1 | 8.648056338 | 9.085807198 | 0.094741997 | 0.43775086 |
| GFRA2 | 8.607944132 | 8.732952106 | 0.707552237 | 0.12500797 |
| GLI1 | 10.25348306 | 10.87518934 | 0.094946771 | 0.62170628 |
| GLI3 | 10.96354145 | 11.61146374 | 0.008579124 | 0.64792229 |
| GPM6A | 9.83485524 | 10.08656501 | 0.399040355 | 0.25170977 |
| GPR135 | 10.55779982 | 11.1994905 | 0.003254375 | 0.64169068 |
| GREB1 | 11.33618173 | 12.6780366 | 0.001204846 | 1.34185487 |
| HHIPL1 | 9.182109522 | 9.701225388 | 0.031879405 | 0.51911587 |
| HOXA4 | 9.336109435 | 9.590652214 | 0.510185537 | 0.25454278 |
| HOXA5 | 8.751398644 | 8.671941493 | 0.800247974 | -0.0794572 |
| IGDCC4 | 10.37270313 | 10.82770854 | 0.03818808 | 0.45500541 |
| KATNAL1 | 11.66410264 | 11.49781818 | 0.299027072 | -0.1662845 |
| KCNMA1 | 11.49092309 | 11.57670601 | 0.742736296 | 0.08578292 |
| KIAA1614 | 10.54661106 | 10.91686287 | 0.067123449 | 0.37025181 |
| KIAA1755 | 10.43462833 | 10.31136484 | 0.573365595 | -0.1232635 |
| KIRREL3 | 8.515556182 | 10.013065 | 1.18E-06 | 1.49750882 |
| LRRC4C | 9.580386127 | 9.938829152 | 0.30083189 | 0.35844302 |
| LRRN2 | 9.331506843 | 9.741575892 | 0.089925115 | 0.41006905 |
| MAGEL2 | 3.582427301 | 7.202176246 | 0.003735283 | 3.61974895 |
| MAGI2 | 10.47820654 | 11.16204267 | 0.001378705 | 0.68383613 |
| MAP1A | 11.57054486 | 11.5661731 | 0.986527867 | -0.0043718 |
| MAP7D3 | 10.82500835 | 10.8806928 | 0.73870332 | 0.05568445 |
| MAP9 | 11.53135874 | 11.48429643 | 0.828744295 | -0.0470623 |
| MDGA1 | 10.87064204 | 11.33133667 | 0.067426134 | 0.46069463 |
| MKX | 9.695408156 | 10.2528398 | 0.004302755 | 0.55743165 |
| MN1 | 11.14015592 | 11.16386713 | 0.928942586 | 0.02371121 |
| MPPED2 | 10.45130907 | 10.49316354 | 0.909021731 | 0.04185447 |
| MYCT1 | 9.686089952 | 9.814993659 | 0.614230487 | 0.12890371 |
| MYOCD | 10.96069186 | 11.60000512 | 0.021107506 | 0.63931325 |
| MYOZ3 | 8.871995956 | 9.64916417 | 0.011224685 | 0.77716821 |
| NACAD | 8.929473492 | 9.611679865 | 0.01290698 | 0.68220637 |
| NFATC4 | 11.13660153 | 11.41818078 | 0.302924058 | 0.28157926 |
| NLGN4X | 10.53701429 | 11.67297051 | 7.89E-06 | 1.13595622 |
| NOVA2 | 10.07495243 | 10.20001186 | 0.609712273 | 0.12505942 |
| NRXN2 | 9.870407271 | 10.62904384 | 0.02955829 | 0.75863657 |
| NUDT10 | 9.426099995 | 9.958975866 | 0.02203288 | 0.53287587 |
| PCDH9 | 10.7162203 | 10.42728497 | 0.301399447 | -0.2889353 |
| PCDHB4 | 9.477477755 | 9.816427337 | 0.125518843 | 0.33894958 |
| PCDHB5 | 8.921065427 | 9.568559332 | 0.001839406 | 0.6474939 |
| PCDHGA3 | 9.121330461 | 9.727190085 | 0.006686441 | 0.60585962 |
| PDE1B | 8.702630063 | 9.531632633 | 0.000274411 | 0.82900257 |
| PDE8B | 10.47229924 | 10.55868936 | 0.606386279 | 0.08639012 |
| PHACTR1 | 10.31897208 | 10.39324551 | 0.748863467 | 0.07427343 |
| PKNOX2 | 9.020377444 | 10.518248 | 8.80E-06 | 1.49787056 |
| PLP1 | 7.077254394 | 7.95031502 | 0.007267772 | 0.87306063 |
| PLXNA4 | 11.11374881 | 11.41921074 | 0.085885944 | 0.30546193 |
| PPFIA2 | 10.04333622 | 10.93714848 | 0.000122708 | 0.89381227 |
| PREX2 | 11.22373169 | 11.26578226 | 0.837989556 | 0.04205057 |
| PRICKLE1 | 10.46099197 | 10.76468861 | 0.287521399 | 0.30369665 |
| PRKD1 | 9.578626293 | 10.22973676 | 0.012608104 | 0.65111047 |
| PRRX1 | 11.42934081 | 11.43360698 | 0.984827466 | 0.00426616 |
| PYGM | 8.657819762 | 8.842222349 | 0.462748537 | 0.18440259 |
| RASGRF2 | 10.36524969 | 11.20135249 | 0.003258416 | 0.83610279 |
| RNF165 | 11.21218641 | 11.67805636 | 0.281268305 | 0.46586995 |
| RUNX1T1 | 12.07412305 | 12.49434662 | 0.03854264 | 0.42022357 |
| SHROOM4 | 10.57375702 | 10.90484935 | 0.086555632 | 0.33109233 |
| SMAD9 | 10.8308871 | 10.73344302 | 0.667435443 | -0.0974441 |
| SMARCD3 | 9.356724819 | 10.5431982 | 1.59E-05 | 1.18647338 |
| SORCS2 | 8.996420104 | 10.34178695 | 0.010073602 | 1.34536685 |
| SSC5D | 11.76170133 | 11.87672656 | 0.477038457 | 0.11502523 |
| SYT15 | 9.65965987 | 10.25840621 | 0.043920981 | 0.59874634 |
| TBC1D9 | 11.16728948 | 11.74101927 | 0.002504322 | 0.5737298 |
| TDRD10 | 7.715724823 | 8.409398866 | 0.013077344 | 0.69367404 |
| TMEM132E | 8.255098897 | 8.413083461 | 0.581204376 | 0.15798456 |
| TNN | 8.837846982 | 9.138572211 | 0.423810626 | 0.30072523 |
| TRIM61 | 7.980789457 | 8.47649125 | 0.035122216 | 0.49570179 |
| TSHZ3 | 10.63300319 | 11.2058644 | 0.050826433 | 0.57286121 |
| TTC7B | 10.47373727 | 11.05028858 | 0.003426462 | 0.57655131 |
| TTLL7 | 11.2003299 | 11.20442308 | 0.986831417 | 0.00409317 |
| UNC5C | 11.22780742 | 11.30941544 | 0.664556301 | 0.08160802 |
| USP51 | 8.932045584 | 9.11021626 | 0.311163032 | 0.17817068 |
| VGLL3 | 11.95019423 | 12.55974316 | 0.006868068 | 0.60954894 |
| WNK3 | 11.34062042 | 12.04937757 | 0.000250093 | 0.70875715 |
| WNT9B | 8.529014837 | 8.6222952 | 0.795355974 | 0.09328036 |
| YPEL4 | 9.093987413 | 9.582087614 | 0.002460781 | 0.4881002 |
| ZFHX4 | 11.84296288 | 12.42753568 | 0.018502539 | 0.5845728 |
| ZFP28 | 10.75733424 | 10.96718801 | 0.290733441 | 0.20985377 |
| ZFPM2 | 10.517483 | 11.28929462 | 0.000195557 | 0.77181162 |
| ZNF135 | 9.793290118 | 10.34552591 | 0.015778389 | 0.55223579 |
| ZNF471 | 11.06642827 | 11.53828544 | 0.06124193 | 0.47185717 |
| ZNF483 | 10.667453 | 10.91333974 | 0.19064778 | 0.24588674 |
| ZNF521 | 10.34407763 | 11.42413515 | 8.97E-06 | 1.08005752 |
| ZNF660 | 10.2725262 | 10.55016195 | 0.060362094 | 0.27763575 |

**Supplementary Table 6. The performance to predict peritoneal or ovarian metastasis by machine learning algorithms and clinical factors.**

| **The expression of msEMT genes to predict Peritoneal/ovarian metastasis** | | | | | | | |
| --- | --- | --- | --- | --- | --- | --- | --- |
| Model | Cohort | Accuracy (95% CI) | NPV | PPV | Specificity | Sensitivity | Delong's test (comarison with TNM) |
| TNM | ACRG (n=300) | 0.798 (0.738-0.858) | 36.8 | 91.5 | 78.1 | 63.6 | Ref |
|  | YCC (n=433) | 0.696 (0.639-0.754) | 27.9 | 91.8 | 83.9 | 45.4 | Ref |
| TNM+Lauren | ACRG (n=292) | 0.827 (0.768-0.885) | 47.9 | 90.9 | 71.4 | 78.6 | 1.17E-06 |
|  | YCC (n=220) | 0.757 (0.688-0.827) | 32 | 94.6 | 89.1 | 50 | 9.61E-05 |
| Elastic Net | ACRG (n=300) | 0.833 (0.786-0.873) | 82.8 | 93.8 | 99.6 | 23.4 | 0.009477 |
|  | YCC (n=433) | 0.811 (0.771-09.847) | 81.6 | 66.7 | 98.6 | 11.5 | 0.0002175 |
| GBM | ACRG (n=300) | 1 (0.988-1) | 100 | 100 | 100 | 100 | 4.20E-11 |
|  | YCC (n=433) | 0.868 (0.816-0.910) | 86.4 | 90.5 | 98.9 | 41.3 | 7.32E-12 |
| SVM | ACRG (n=300) | 0.887 (0.845-0.920) | 88.3 | 91.7 | 98.7 | 51.6 | 4.81E-07 |
|  | YCC (n=433) | 0.892 (0.858-0.919) | 89.7 | 85.7 | 97.7 | 55.2 | 2.20E-16 |
| RF | ACRG (n=300) | 1 (0.988-1) | 100 | 100 | 100 | 100 | 4.20E-11 |
|  | YCC (n=433) | 1 (0.992-1) | 100 | 100 | 100 | 100 | 2.20E-16 |
| PCANN | ACRG (n=300) | 0.787 (0.736-0.831) | 78.7 | NA | 100 | 0 | 9.42E-03 |
|  | YCC (n=433) | 0.799 (0.758-0.836) | 79.9 | NA | 100 | 0 | 1.63E-11 |
| **The expression of msEMT genes with TNM and Lauren model to predict peritoneal/ovarian metstasis** | | | | | | | |
| Model*  (TNM + Lauren) | Cohort | Accuracy (95% CI) | NPV | PPV | Specificity | Sensitivity | Delong's test (comarison with TNM) |
| Elastic Net | ACRG (n=292) | 0.88 (0.837-0.915) | 87.6 | 91.2 | 98.7 | 49.2 | 2.20E-16 |
|  | YCC (n=220) | 0.82 (0.761-0.867) | 83.2 | 66.7 | 96.6 | 26.1 | 4.23E-09 |
| GBM | ACRG (n=292) | 0.849 (0.803-0.888) | 84.1 | 95.2 | 99.6 | 31.8 | 2.20E-16 |
|  | YCC (n=220) | 0.932 (0.89-0.961) | 93 | 94.3 | 98.9 | 71.7 | 2.20E-16 |
| SVM | ACRG (n=292) | 0.938 (0.904-0.963) | 93.8 | 94.1 | 98.7 | 76.2 | 2.20E-16 |
|  | YCC (n=220) | 0.932 (0.89-0.961) | 92.5 | 97 | 99.4 | 69.6 | 2.20E-16 |
| RF | ACRG (n=292) | 1 (0.987-1) | 100 | 100 | 100 | 100 | 2.20E-16 |
|  | YCC (n=220) | 1 (0.983-1) | 100 | 100 | 100 | 100 | 2.20E-16 |
| PCANN | ACRG (n=292) | 0.784 (0.733-0.83) | 78.4 | NA | 100 | 0 | 8.60E-02 |
|  | YCC (n=220) | 0.791 (0.731-0.843) | 79.1 | NA | 100 | 0 | 5.397E-07 |

*The models includes TNM + Lauren.

**Supplementary Table 7. The origin of gastric cancer cell lines and the log2 expression of msEMT genes in each cell line.**

| **GC_cell-line_id** | **Tumor_origin** |  | Symbol | AGS | MKN28 | MKN45 | MKN74 | NCC20 | NCC24 | NCC59 | SNU1 | SNU16 | SNU1750 | SNU484 | SNU601 | SNU620 | SNU638 | SNU668 | SNU719 | YCC11 | YCC2 | YCC3 |
| --- | --- | --- | --- | --- | --- | --- | --- | --- | --- | --- | --- | --- | --- | --- | --- | --- | --- | --- | --- | --- | --- | --- |
| AGS | Primary |  | FMO1 | 0 | 1.5849625 | 1 | 0 | 0 | 1.5849625 | 2 | 0 | 0 | 0 | 0 | 0 | 0 | 0 | 0 | 0 | 0 | 0 | 0 |
| MKN28 | Liver |  | PREX2 | 0 | 0 | 1.5849625 | 0 | 1 | 0 | 0 | 1 | 1.5849625 | 9.25502857 | 3.5849625 | 0 | 1.5849625 | 0 | 0 | 0 | 1 | 1 | 0 |
| MKN45 | Liver |  | CDON | 4.39231742 | 6.12928302 | 7.91288934 | 5.78135971 | 5.78135971 | 6.83289001 | 8.8917837 | 10.509775 | 6.08746284 | 7.67242534 | 5.72792045 | 6.74146699 | 6.59991284 | 7.28540222 | 6.22881869 | 6.68650053 | 8.56605404 | 8.15481811 | 6.64385619 |
| MKN74 | Liver |  | MPPED2 | 0 | 1 | 2 | 0 | 0 | 1 | 1.5849625 | 1 | 1.5849625 | 5.9068906 | 1.5849625 | 0 | 1.5849625 | 1 | 1.5849625 | 0 | 8.39231742 | 0 | 0 |
| NCC20 | Ascites |  | PYGM | 2.5849625 | 4.52356196 | 2.80735492 | 4.08746284 | 4.32192809 | 3.5849625 | 4.95419631 | 2.5849625 | 2 | 5.55458885 | 2.32192809 | 1.5849625 | 3.169925 | 1.5849625 | 2.32192809 | 3.5849625 | 3.70043972 | 2.32192809 | 2.80735492 |
| NCC24 | Primary |  | ABCC9 | 7.48381578 | 10.4008794 | 8.91587938 | 10.504819 | 8.16490693 | 6.857981 | 8.63662462 | 7.28540222 | 7.08746284 | 10.0402897 | 8.15987134 | 8.37503943 | 7.54689446 | 7.33985 | 8.20457114 | 7.94836723 | 8.32642949 | 8.56605404 | 8.6329952 |
| NCC59 | Ascites |  | RUNX1T1 | 1 | 0 | 2.80735492 | 0 | 0 | 0 | 0 | 0 | 0 | 3.45943162 | 4.52356196 | 0 | 0 | 0 | 0 | 0 | 0 | 0 | 0 |
| SNU1 | Primary |  | PCDHB4 | 2.5849625 | 2.32192809 | 4.08746284 | 2 | 1 | 3 | 1 | 0 | 1 | 2.32192809 | 0 | 0 | 1 | 1 | 0 | 3.45943162 | 2.5849625 | 1.5849625 | 3.70043972 |
| SNU1750 | Primary |  | SMARCD3 | 8.74146699 | 2.5849625 | 7.74146699 | 5.88264305 | 5.08746284 | 6.76818432 | 7.53915881 | 10.7202443 | 5.169925 | 8.34872815 | 10.4604559 | 7.31288296 | 7.9248125 | 9.75321675 | 8.68999797 | 4.80735492 | 9.32642949 | 7.0768156 | 7.94251451 |
| SNU484 | Primary |  | ZFHX4 | 0 | 1 | 1 | 1 | 5.357552 | 2.32192809 | 3.32192809 | 0 | 4.80735492 | 6 | 10.1522848 | 0 | 1 | 2.32192809 | 4.08746284 | 0 | 8.88874325 | 0 | 3 |
| SNU601 | Ascites |  | NFATC4 | 4.169925 | 4.5849625 | 8.34429591 | 6.26678654 | 7.0768156 | 4.70043972 | 2 | 0 | 8.49984589 | 9.05799172 | 7.93663794 | 4.32192809 | 4.80735492 | 7.11894107 | 2.80735492 | 4.169925 | 1.5849625 | 6.20945337 | 3.45943162 |
| SNU638 | Ascites |  | KATNAL1 | 8.82017896 | 8.95128471 | 4.39231742 | 8.53915881 | 2.32192809 | 4 | 7.27612441 | 9.14974712 | 2.80735492 | 8.92777796 | 10.082149 | 6.47573343 | 7.56224242 | 8.23840474 | 9.45532722 | 6.169925 | 8.40939094 | 8.08214904 | 6.5849625 |
| SNU668 | Ascites |  | CORO2B | 0 | 0 | 2.5849625 | 5.169925 | 0 | 0 | 0 | 0 | 0 | 6.04439412 | 8.69696753 | 0 | 0 | 0 | 3 | 0 | 1 | 5.169925 | 2.32192809 |
| SNU719 | Primary |  | IGDCC4 | 2 | 6.76818432 | 6.04439412 | 6.37503943 | 4.52356196 | 2 | 2.32192809 | 3.169925 | 2.32192809 | 6.84549005 | 6.62935662 | 0 | 0 | 3.9068906 | 5.83289001 | 2 | 4 | 6.12928302 | 7.04439412 |
| YCC11 | Ascites |  | NOVA2 | 0 | 4 | 0 | 3.70043972 | 3 | 0 | 4.32192809 | 2.80735492 | 0 | 2.5849625 | 4.52356196 | 3 | 3.9068906 | 2.80735492 | 2 | 0 | 3.32192809 | 3.32192809 | 3.169925 |
| YCC2 | Ascites |  | HOXA5 | 0 | 4.52356196 | 6.80735492 | 4.70043972 | 5.70043972 | 0 | 7.47573343 | 6.53915881 | 6.61470984 | 3.5849625 | 6.98868469 | 0 | 0 | 8.28540222 | 5.24792751 | 7.357552 | 9.00562455 | 5.95419631 | 6.68650053 |
| YCC3 | Ascites |  | GLI3 | 1 | 2.32192809 | 0 | 8.23361968 | 0 | 1.5849625 | 0 | 9.99859043 | 2 | 11.2312212 | 1 | 0 | 0 | 0 | 10.6617781 | 3.70043972 | 5.24792751 | 8.40087944 | 2 |
| SNU16 | Ascites |  | EBF3 | 1 | 0 | 4.5849625 | 0 | 1 | 2.5849625 | 2.32192809 | 7.4918531 | 0 | 3 | 0 | 0 | 0 | 0 | 0 | 0 | 0 | 0 | 0 |
| SNU620 | Ascites |  | TBC1D9 | 4.39231742 | 1.5849625 | 7.23840474 | 1 | 4.95419631 | 7.87651695 | 6.83289001 | 0 | 8.38370429 | 9.08746284 | 8.57364719 | 1.5849625 | 0 | 7.53138146 | 8.24317398 | 1 | 10.2691267 | 8 | 10.0768156 |
|  |  |  | NRXN2 | 1.5849625 | 2 | 3 | 4.169925 | 5 | 3.9068906 | 0 | 1.5849625 | 0 | 4.39231742 | 2.32192809 | 1 | 3.70043972 | 0 | 2.80735492 | 1 | 4.857981 | 4.80735492 | 5.12928302 |
|  |  |  | GLI1 | 5.12928302 | 0 | 2.80735492 | 4.857981 | 0 | 4 | 7.10852446 | 8.9248125 | 6.06608919 | 3.32192809 | 8.30833903 | 3 | 2.5849625 | 6.42626475 | 2.80735492 | 4 | 6.88264305 | 3.45943162 | 5.55458885 |
|  |  |  | PHACTR1 | 2.5849625 | 2.80735492 | 3.5849625 | 1.5849625 | 2.32192809 | 2.32192809 | 0 | 2.80735492 | 0 | 4.7548875 | 2.5849625 | 3.32192809 | 2.32192809 | 2 | 0 | 3.169925 | 0 | 2.32192809 | 2.32192809 |
|  |  |  | MDGA1 | 0 | 3.70043972 | 5.5849625 | 3.9068906 | 0 | 4.24792751 | 5.95419631 | 1 | 0 | 2.32192809 | 7.11894107 | 3.32192809 | 1.5849625 | 5.08746284 | 6.4918531 | 5 | 5.4918531 | 5.4918531 | 6.82017896 |
|  |  |  | EPM2A | 7.02236781 | 4.169925 | 4.39231742 | 6.55458885 | 5.169925 | 5.7548875 | 7.56224242 | 4.5849625 | 6.37503943 | 6.78135971 | 5.70043972 | 5.12928302 | 6.14974712 | 6.24792751 | 5.20945337 | 7.05528244 | 6.64385619 | 4.70043972 | 5.64385619 |
|  |  |  | PCDHB5 | 2 | 1.5849625 | 1 | 1.5849625 | 0 | 9.18487534 | 1 | 0 | 1 | 10.8933015 | 6.71424552 | 1 | 0 | 2 | 2.5849625 | 0 | 8.04439412 | 2.32192809 | 8.17492568 |
|  |  |  | PDE8B | 4.80735492 | 4.52356196 | 5.4918531 | 4.64385619 | 4.08746284 | 4.24792751 | 4.70043972 | 4.7548875 | 4.39231742 | 6.24792751 | 5.12928302 | 5.08746284 | 3.32192809 | 4.64385619 | 10.4938554 | 4.32192809 | 9.79928162 | 6.88264305 | 5.80735492 |
|  |  |  | RASGRF2 | 5 | 3.169925 | 2.80735492 | 2.5849625 | 3.80735492 | 1.5849625 | 4.7548875 | 3.5849625 | 1 | 5.61470984 | 8.80413102 | 2.32192809 | 2 | 5.357552 | 4 | 2.32192809 | 2.80735492 | 3.9068906 | 1.5849625 |
|  |  |  | BCHE | 0 | 0 | 0 | 0 | 0 | 1.5849625 | 4.169925 | 0 | 3.5849625 | 2 | 1.5849625 | 1.5849625 | 0 | 0 | 3.80735492 | 2 | 9.95274125 | 3.45943162 | 0 |
|  |  |  | PRRX1 | 0 | 0 | 0 | 0 | 0 | 0 | 1 | 0 | 0 | 6.64385619 | 4.64385619 | 0 | 0 | 0 | 0 | 0 | 3.45943162 | 0 | 0 |
|  |  |  | MYCT1 | 0 | 0 | 0 | 2 | 0 | 1 | 1.5849625 | 0 | 0 | 1.5849625 | 3.169925 | 0 | 0 | 2.32192809 | 0 | 2.5849625 | 0 | 1.5849625 | 0 |
|  |  |  | TNN | 1 | 1 | 1 | 0 | 0 | 0 | 0 | 0 | 0 | 7.31288296 | 0 | 0 | 0 | 0 | 0 | 0 | 0 | 0 | 0 |
|  |  |  | SMAD9 | 1.5849625 | 3 | 1.5849625 | 5.08746284 | 2.32192809 | 0 | 5 | 5.55458885 | 8.41362793 | 8.07146236 | 7.70735913 | 1 | 3.32192809 | 1 | 7.45121111 | 1 | 6.30378075 | 1.5849625 | 1 |
|  |  |  | TSHZ3 | 0 | 0 | 5.61470984 | 5.83289001 | 0 | 2.32192809 | 0 | 0 | 0 | 3.32192809 | 3.80735492 | 0 | 0 | 0 | 0 | 0 | 0 | 0 | 4.24792751 |
|  |  |  | NUDT10 | 0 | 0 | 0 | 0 | 0 | 0 | 0 | 0 | 0 | 9.91438513 | 5.20945337 | 0 | 0 | 0 | 1 | 0 | 0 | 1 | 0 |
|  |  |  | PDE1B | 0 | 1 | 7.09803208 | 0 | 1 | 1 | 2.5849625 | 0 | 0 | 2.32192809 | 1 | 0 | 0 | 0 | 0 | 3.5849625 | 3 | 0 | 2.32192809 |
|  |  |  | PLP1 | 4.169925 | 0 | 0 | 2 | 0 | 0 | 0 | 0 | 0 | 2.80735492 | 1 | 2.5849625 | 0 | 0 | 0 | 0 | 0 | 0 | 0 |
|  |  |  | FAM124B | 2 | 0 | 1 | 1 | 0 | 1.5849625 | 1 | 0 | 0 | 1.5849625 | 0 | 1 | 0 | 0 | 0 | 0 | 1 | 1 | 0 |
|  |  |  | CCDC136 | 4.64385619 | 2 | 1.5849625 | 4.52356196 | 4.64385619 | 3.9068906 | 6.14974712 | 2.32192809 | 2.80735492 | 7.09803208 | 6.39231742 | 0 | 1 | 6.72792045 | 7.169925 | 0 | 5.24792751 | 4.80735492 | 5.4918531 |
|  |  |  | ARHGAP22 | 3.5849625 | 4.169925 | 5.88264305 | 4.70043972 | 5.12928302 | 4.169925 | 5.70043972 | 4 | 4.24792751 | 5.5849625 | 6.10852446 | 4.70043972 | 4.52356196 | 5 | 4.857981 | 4.08746284 | 4.7548875 | 5.24792751 | 4.39231742 |
|  |  |  | ATP1B2 | 0 | 2.32192809 | 1.5849625 | 3.70043972 | 0 | 1 | 1 | 2 | 2.5849625 | 8.48381578 | 3 | 2.32192809 | 1.5849625 | 0 | 3.32192809 | 1 | 9.13955135 | 0 | 3.45943162 |
|  |  |  | MAP7D3 | 7.87036472 | 4.24792751 | 8.34429591 | 4.64385619 | 7.02236781 | 8.33985 | 9.22881869 | 10.3586512 | 8.84235034 | 9.16239133 | 10.7540524 | 7.42626475 | 4.08746284 | 8.91587938 | 9.38154295 | 5.20945337 | 9.81698362 | 9.19229281 | 8.92777796 |
|  |  |  | EDA2R | 2 | 0 | 6.4918531 | 2.80735492 | 5.39231742 | 1 | 2 | 0 | 1 | 2.80735492 | 3.169925 | 0 | 0 | 0 | 5.28540222 | 0 | 1 | 0 | 0 |
|  |  |  | DLG4 | 5.32192809 | 5.45943162 | 9.57364719 | 6.79441587 | 3.169925 | 5.39231742 | 4.52356196 | 7.78790256 | 2.32192809 | 8.53915881 | 9.72962074 | 6.96578428 | 4.24792751 | 6.37503943 | 8.40939094 | 3.9068906 | 9.20701432 | 7.76155123 | 4.24792751 |
|  |  |  | FHOD3 | 0 | 3 | 2.80735492 | 5.5849625 | 4.52356196 | 3.45943162 | 4.169925 | 4.95419631 | 3.169925 | 9.29462075 | 6 | 0 | 0 | 3.70043972 | 8.14465824 | 2.32192809 | 8.04984855 | 8.09803208 | 2.80735492 |
|  |  |  | ANKRD6 | 0 | 7.41785251 | 6 | 4.52356196 | 2 | 4.24792751 | 1.5849625 | 1.5849625 | 2 | 7.83289001 | 7.32192809 | 1 | 1.5849625 | 1 | 7.11894107 | 1 | 5.08746284 | 6.7548875 | 7.36632221 |
|  |  |  | KIAA1614 | 3.45943162 | 6.22881869 | 3.9068906 | 4 | 6.65821148 | 2 | 3.80735492 | 5.64385619 | 3.5849625 | 7.47573343 | 6.14974712 | 5.70043972 | 2.80735492 | 5.45943162 | 6.04439412 | 2.5849625 | 7.18982456 | 4.52356196 | 4.64385619 |
|  |  |  | NACAD | 2.80735492 | 3.169925 | 1 | 5.28540222 | 6.22881869 | 0 | 3.45943162 | 1 | 1 | 8.56605404 | 6.98868469 | 2.32192809 | 0 | 7.29462075 | 6.4429435 | 0 | 4.7548875 | 5.67242534 | 1 |
|  |  |  | ARHGAP20 | 0 | 1 | 0 | 0 | 0 | 2.5849625 | 0 | 0 | 1 | 6.40939094 | 1.5849625 | 0 | 0 | 0 | 0 | 0 | 4.64385619 | 0 | 0 |
|  |  |  | TTLL7 | 5.12928302 | 4.857981 | 3.70043972 | 4.52356196 | 4.24792751 | 4.5849625 | 8.08746284 | 4.9068906 | 5.08746284 | 7.68650053 | 8.57364719 | 4.5849625 | 4.24792751 | 6.12928302 | 4.70043972 | 3.80735492 | 7.85174904 | 2.32192809 | 7.54689446 |
|  |  |  | PRICKLE1 | 0 | 7.4429435 | 0 | 8.36194377 | 1 | 0 | 0 | 0 | 0 | 8.43879185 | 8.57364719 | 0 | 0 | 4.64385619 | 5.28540222 | 2.80735492 | 2.5849625 | 7.18982456 | 6.4918531 |
|  |  |  | PPFIA2 | 0 | 0 | 0 | 0 | 1 | 2.5849625 | 1 | 2 | 0 | 2 | 4.39231742 | 0 | 0 | 0 | 2.5849625 | 0 | 0 | 0 | 0 |
|  |  |  | MYOCD | 0 | 0 | 0 | 0 | 0 | 0 | 0 | 0 | 0 | 2 | 2.5849625 | 0 | 0 | 0 | 1 | 0 | 11.8185822 | 0 | 2.5849625 |
|  |  |  | RNF165 | 2.32192809 | 2.32192809 | 5.45943162 | 2.5849625 | 7.21916852 | 2 | 3.70043972 | 2.80735492 | 2.5849625 | 5.08746284 | 3.9068906 | 1.5849625 | 1.5849625 | 2 | 3 | 3.80735492 | 6.52356196 | 0 | 2.32192809 |
|  |  |  | CASQ1 | 0 | 0 | 0 | 1 | 2.80735492 | 1.5849625 | 1.5849625 | 1.5849625 | 2 | 5.67242534 | 1 | 2 | 0 | 0 | 0 | 0 | 4.24792751 | 0 | 1 |
|  |  |  | ANKRD53 | 0 | 1 | 3.5849625 | 1 | 1.5849625 | 1 | 1 | 1 | 0 | 4.24792751 | 0 | 1 | 1 | 1 | 2.32192809 | 0 | 0 | 0 | 1.5849625 |
|  |  |  | CAND2 | 1 | 3.169925 | 3.45943162 | 2.5849625 | 3 | 3 | 2.80735492 | 3.169925 | 3.5849625 | 7.98299357 | 8.97441459 | 2.5849625 | 2.32192809 | 1.5849625 | 5.88264305 | 0 | 1.5849625 | 4.08746284 | 3.80735492 |
|  |  |  | ZNF660 | 2 | 2.5849625 | 3 | 1 | 2.80735492 | 2 | 1.5849625 | 4.08746284 | 1.5849625 | 7.22881869 | 2.80735492 | 1 | 3.70043972 | 2.80735492 | 2.32192809 | 2 | 2.32192809 | 3.45943162 | 1 |
|  |  |  | ADPRH | 0 | 7.22881869 | 6.14974712 | 7.29462075 | 5.67242534 | 0 | 0 | 0 | 2.80735492 | 6.93073734 | 5.5849625 | 0 | 0 | 6.02236781 | 4.45943162 | 0 | 3.70043972 | 5.9068906 | 4.32192809 |
|  |  |  | NLGN4X | 0 | 1 | 0 | 0 | 0 | 2.5849625 | 0 | 0 | 0 | 9.81858218 | 9.20701432 | 0 | 0 | 0 | 0 | 0 | 0 | 0 | 0 |
|  |  |  | FAM13C | 0 | 0 | 0 | 1.5849625 | 1 | 0 | 1 | 0 | 0 | 2 | 5 | 0 | 0 | 2.32192809 | 0 | 0 | 3.45943162 | 0 | 0 |
|  |  |  | LRRC4C | 1 | 0 | 0 | 0 | 5.12928302 | 0 | 0 | 0 | 0 | 4.32192809 | 3.169925 | 0 | 0 | 2.80735492 | 0 | 0 | 0 | 0 | 1 |
|  |  |  | KIRREL3 | 1 | 4.64385619 | 2.5849625 | 3.80735492 | 4 | 2.80735492 | 2.80735492 | 7.42626475 | 3.5849625 | 6.10852446 | 3.70043972 | 2.5849625 | 1 | 0 | 3.5849625 | 1.5849625 | 4.32192809 | 3.32192809 | 1 |
|  |  |  | KIAA1755 | 1 | 4.24792751 | 0 | 5.42626475 | 0 | 1.5849625 | 1.5849625 | 3.70043972 | 0 | 2.80735492 | 1.5849625 | 2 | 0 | 1.5849625 | 2 | 1 | 0 | 3.32192809 | 0 |
|  |  |  | CNKSR2 | 0 | 1 | 0 | 0 | 2 | 1 | 0 | 0 | 0 | 7.59991284 | 5.5849625 | 0 | 0 | 0 | 5.04439412 | 0 | 0 | 3 | 0 |
|  |  |  | MKX | 0 | 0 | 7.033423 | 0 | 0 | 0 | 0 | 0 | 3 | 5.55458885 | 3.80735492 | 0 | 0 | 1 | 1.5849625 | 0 | 3.9068906 | 1.5849625 | 3.169925 |
|  |  |  | FAM124A | 2.80735492 | 1.5849625 | 1.5849625 | 4.64385619 | 2 | 5.04439412 | 4.52356196 | 6.45943162 | 2 | 4.08746284 | 5.169925 | 1 | 3.9068906 | 6.84549005 | 3.45943162 | 2.80735492 | 7.47573343 | 2 | 3 |
|  |  |  | GPM6A | 0 | 0 | 0 | 2 | 0 | 5.55458885 | 0 | 0 | 0 | 6.72792045 | 5.32192809 | 0 | 0 | 3.70043972 | 0 | 0 | 4 | 2 | 0 |
|  |  |  | DLG2 | 2.5849625 | 4.24792751 | 4.45943162 | 4.45943162 | 2.32192809 | 2.32192809 | 4 | 3.5849625 | 2.5849625 | 3.70043972 | 4.80735492 | 2.5849625 | 2.32192809 | 5.357552 | 7.27612441 | 4 | 4.169925 | 4.95419631 | 2.32192809 |
|  |  |  | DIXDC1 | 6.39231742 | 5.357552 | 7.41785251 | 6.67242534 | 6.4429435 | 7.46760555 | 4.39231742 | 8.74819285 | 8.14465824 | 7.91288934 | 9.36850646 | 5.93073734 | 7.22881869 | 7.33985 | 8.47573343 | 7.24792751 | 9.25029842 | 6.97727992 | 6.169925 |
|  |  |  | AKAP6 | 3.70043972 | 4.5849625 | 5.08746284 | 5.42626475 | 3 | 4.64385619 | 8.62935662 | 2.80735492 | 5.70043972 | 7.62205182 | 3 | 4.52356196 | 4.95419631 | 5.97727992 | 3 | 6.80735492 | 4.5849625 | 5.45943162 | 6.12928302 |
|  |  |  | CACNA2D1 | 0 | 4.5849625 | 0 | 0 | 0 | 4.169925 | 8.16490693 | 0 | 4.169925 | 6.37503943 | 11.693487 | 0 | 1 | 7 | 3 | 3.9068906 | 8.76487159 | 1 | 0 |
|  |  |  | FGD5 | 2.32192809 | 3 | 4.08746284 | 3.9068906 | 4.52356196 | 4.39231742 | 3.169925 | 3.80735492 | 2.5849625 | 4.7548875 | 4.9068906 | 3.9068906 | 3.9068906 | 3.5849625 | 3.169925 | 3.5849625 | 4 | 3.169925 | 3.5849625 |
|  |  |  | KCNMA1 | 1.5849625 | 0 | 2.5849625 | 1.5849625 | 0 | 2.5849625 | 2.32192809 | 3 | 7 | 12.3459596 | 8.49585503 | 0 | 0 | 0 | 6.30378075 | 0 | 1.5849625 | 7.15987134 | 1 |
|  |  |  | ADAMTSL3 | 6.06608919 | 8.857981 | 7.88874325 | 8.87036472 | 7.41785251 | 5.42626475 | 8.11894107 | 7.31288296 | 5.169925 | 7.11894107 | 7.5849625 | 7.02236781 | 7.12928302 | 5.5849625 | 6.80735492 | 6.68650053 | 7.0768156 | 7.88874325 | 6.79441587 |
|  |  |  | ERG | 0 | 0 | 0 | 0 | 0 | 2.32192809 | 0 | 0 | 0 | 0 | 0 | 0 | 0 | 0 | 4.64385619 | 0 | 0 | 0 | 2.32192809 |
|  |  |  | SHROOM4 | 3 | 3.32192809 | 4.169925 | 0 | 1.5849625 | 2.32192809 | 0 | 5.04439412 | 2.32192809 | 4.32192809 | 2.32192809 | 2 | 0 | 3 | 3.9068906 | 3 | 3.169925 | 4 | 2.32192809 |
|  |  |  | DYNC1I1 | 3 | 6.67242534 | 6.26678654 | 7.04439412 | 0 | 2 | 2.32192809 | 4.24792751 | 0 | 9.94690627 | 4 | 2 | 3 | 2 | 3.9068906 | 6.45943162 | 6.79441587 | 3.169925 | 0 |
|  |  |  | WNT9B | 2.5849625 | 0 | 0 | 1 | 3.169925 | 1 | 2.5849625 | 2.80735492 | 2 | 2.32192809 | 2.80735492 | 1 | 1.5849625 | 2 | 3 | 1.5849625 | 2 | 0 | 0 |
|  |  |  | C1QTNF7 | 0 | 0 | 0 | 0 | 0 | 0 | 0 | 0 | 0 | 0 | 0 | 0 | 0 | 1.5849625 | 0 | 0 | 0 | 0 | 0 |
|  |  |  | TDRD10 | 0 | 0 | 0 | 2 | 0 | 2.32192809 | 0 | 0 | 0 | 2.32192809 | 1 | 0 | 0 | 0 | 1.5849625 | 0 | 1.5849625 | 0 | 1 |
|  |  |  | MAP9 | 0 | 0 | 7.18982456 | 0 | 8.54303182 | 0 | 0 | 0 | 0 | 8.91288934 | 7.91886324 | 0 | 0 | 0 | 8.08746284 | 0 | 7.96000193 | 0 | 4.32192809 |
|  |  |  | CMYA5 | 2.32192809 | 6.18982456 | 5.857981 | 6.32192809 | 2.5849625 | 2.32192809 | 5.42626475 | 4.95419631 | 3.32192809 | 4.64385619 | 2.32192809 | 2.32192809 | 2.32192809 | 5.32192809 | 3 | 4.24792751 | 4.5849625 | 4.95419631 | 4.80735492 |
|  |  |  | MYOZ3 | 1 | 1 | 1.5849625 | 2.32192809 | 0 | 3.32192809 | 1 | 0 | 3 | 7.99435344 | 5.357552 | 1.5849625 | 0 | 1 | 2.5849625 | 1 | 1 | 4.32192809 | 1 |
|  |  |  | PKNOX2 | 0 | 3.45943162 | 0 | 0 | 0 | 1.5849625 | 1.5849625 | 6.08746284 | 0 | 4.857981 | 0 | 1 | 0 | 0 | 0 | 0 | 0 | 1.5849625 | 0 |
|  |  |  | TTC7B | 7.63662462 | 7.55458885 | 7 | 8.50382574 | 3.45943162 | 0 | 3.32192809 | 9.42626475 | 8.81057163 | 9.03617361 | 9.81698362 | 3 | 2 | 8.91886324 | 6.78135971 | 0 | 7.30378075 | 9.09803208 | 8.11374217 |
|  |  |  | YPEL4 | 1.5849625 | 2.80735492 | 0 | 2 | 0 | 0 | 0 | 1 | 0 | 4 | 2.5849625 | 0 | 0 | 1.5849625 | 0 | 1 | 4.169925 | 1.5849625 | 2 |
|  |  |  | MAP1A | 4.857981 | 4.39231742 | 5.4918531 | 6.94251451 | 4.32192809 | 2.32192809 | 2.5849625 | 3.70043972 | 3 | 10.3695973 | 8.42206477 | 3.70043972 | 4.24792751 | 7.5849625 | 9.25974326 | 4.64385619 | 9.64385619 | 4.24792751 | 3.70043972 |
|  |  |  | GFRA2 | 0 | 0 | 1 | 0 | 3 | 0 | 0 | 0 | 0 | 7.37503943 | 0 | 0 | 0 | 0 | 0 | 0 | 1.5849625 | 0 | 0 |
|  |  |  | FAM110B | 8.77148947 | 0 | 0 | 0 | 4.24792751 | 9.49785184 | 1.5849625 | 0 | 0 | 5.169925 | 3 | 0 | 9.41996018 | 0 | 1 | 0 | 6.79441587 | 2.80735492 | 8.38801729 |
|  |  |  | CPT1C | 7.30378075 | 4.24792751 | 3.70043972 | 7.52356196 | 4.45943162 | 5.45943162 | 2.5849625 | 1.5849625 | 5.32192809 | 10.0402897 | 11.2390018 | 5.88264305 | 3.80735492 | 7.22881869 | 4.70043972 | 2.5849625 | 9.67771964 | 1 | 5.39231742 |
|  |  |  | MN1 | 0 | 2 | 2.5849625 | 4.9068906 | 0 | 0 | 2.5849625 | 6.53915881 | 0 | 8.75154406 | 4.64385619 | 4.08746284 | 0 | 0 | 7.28540222 | 1 | 7.857981 | 9.5372184 | 4.70043972 |
|  |  |  | ZFPM2 | 3.32192809 | 4.95419631 | 4.24792751 | 4.9068906 | 2.32192809 | 3.169925 | 4.32192809 | 5 | 3 | 7.91886324 | 8.30833903 | 2 | 1.5849625 | 1 | 3.5849625 | 4.39231742 | 4 | 3.9068906 | 4.169925 |
|  |  |  | LRRN2 | 1 | 0 | 6.24792751 | 3 | 2 | 1 | 2.5849625 | 2.5849625 | 3.70043972 | 5.28540222 | 7.25738784 | 1 | 1 | 1 | 7.10852446 | 0 | 3.9068906 | 8.33539035 | 6.4429435 |
|  |  |  | DSEL | 1.5849625 | 3 | 3 | 3.70043972 | 2.5849625 | 2 | 1.5849625 | 2.32192809 | 2.32192809 | 7.41785251 | 8.10328781 | 1 | 0 | 2 | 9.9248125 | 1 | 9.86573327 | 1.5849625 | 4 |
|  |  |  | ZNF483 | 2.80735492 | 2.5849625 | 3.45943162 | 4.39231742 | 1.5849625 | 2.80735492 | 3 | 6.65821148 | 3.169925 | 5.24792751 | 5.93073734 | 3.9068906 | 4.169925 | 5.357552 | 2 | 3.169925 | 6.67242534 | 4.45943162 | 3.32192809 |
|  |  |  | ZNF135 | 0 | 0 | 0 | 0 | 0 | 0 | 0 | 0 | 0 | 0 | 1 | 0 | 0 | 0 | 0 | 0 | 1 | 0 | 0 |
|  |  |  | CCDC89 | 1.5849625 | 1 | 0 | 1.5849625 | 1 | 1 | 0 | 1 | 1 | 2.32192809 | 1 | 1.5849625 | 0 | 6.74146699 | 0 | 0 | 4.64385619 | 3.70043972 | 0 |
|  |  |  | SSC5D | 2.80735492 | 4.857981 | 3.32192809 | 2 | 2.32192809 | 1 | 3 | 5.55458885 | 0 | 4.5849625 | 4.9068906 | 2.80735492 | 1.5849625 | 1 | 3.70043972 | 4.70043972 | 3.80735492 | 1.5849625 | 1.5849625 |
|  |  |  | TMEM132E | 0 | 0 | 0 | 0 | 2 | 2.32192809 | 0 | 0 | 0 | 5.04439412 | 5.20945337 | 0 | 0 | 0 | 1.5849625 | 0 | 0 | 0 | 4 |
|  |  |  | GPR135 | 2.80735492 | 2.5849625 | 3.169925 | 3.169925 | 3.32192809 | 3.5849625 | 3.45943162 | 4.52356196 | 4.08746284 | 4.70043972 | 6.47573343 | 3.169925 | 4.70043972 | 4.24792751 | 6.40939094 | 3.70043972 | 5.97727992 | 6.10852446 | 3.80735492 |
|  |  |  | UNC5C | 1 | 1 | 0 | 0 | 0 | 0 | 3.32192809 | 0 | 2 | 5.04439412 | 2.80735492 | 0 | 0 | 0 | 0 | 0 | 6.84549005 | 0 | 1 |
|  |  |  | HHIPL1 | 0 | 0 | 2 | 1 | 1.5849625 | 2 | 2 | 0 | 0 | 5.5849625 | 7.26678654 | 1 | 0 | 1 | 4.64385619 | 2 | 2.32192809 | 7.65105169 | 1.5849625 |
|  |  |  | TRIM61 | 0 | 1.5849625 | 1 | 2 | 0 | 1 | 0 | 0 | 3 | 0 | 0 | 0 | 2 | 0 | 2 | 0 | 1 | 1 | 0 |
|  |  |  | FBXL7 | 1 | 0 | 0 | 0 | 0 | 2 | 0 | 1 | 0 | 7.97154355 | 8.4918531 | 1 | 0 | 0 | 4 | 0 | 4 | 0 | 1 |
|  |  |  | PCDH9 | 0 | 0 | 0 | 0 | 0 | 1 | 1 | 1 | 3 | 6.97727992 | 8.74483384 | 0 | 0 | 0 | 8.24317398 | 0 | 6.78135971 | 4.169925 | 0 |
|  |  |  | PRKD1 | 0 | 0 | 6.08746284 | 0 | 0 | 0 | 2.5849625 | 5.61470984 | 0 | 6.40939094 | 8.70390357 | 0 | 3 | 0 | 3.32192809 | 0 | 1 | 3.169925 | 2.5849625 |
|  |  |  | SORCS2 | 1.5849625 | 0 | 1 | 1 | 1.5849625 | 1 | 2 | 1.5849625 | 0 | 2 | 3.70043972 | 1 | 0 | 1 | 3.32192809 | 0 | 3 | 8.70043972 | 8.08746284 |
|  |  |  | C3orf70 | 0 | 2 | 2.80735492 | 2.5849625 | 3.45943162 | 1 | 2.5849625 | 3.9068906 | 3.9068906 | 3.169925 | 2.5849625 | 2 | 2.32192809 | 3.169925 | 1.5849625 | 0 | 3 | 1 | 4 |
|  |  |  | MAGI2 | 0 | 3.70043972 | 6.65821148 | 4.32192809 | 4.24792751 | 0 | 1 | 0 | 5.28540222 | 4.32192809 | 7.27612441 | 0 | 1.5849625 | 3 | 6.32192809 | 0 | 4.9068906 | 4.7548875 | 3.80735492 |
|  |  |  | GREB1 | 3.5849625 | 5.04439412 | 5.169925 | 6.12928302 | 5.20945337 | 5.28540222 | 4.52356196 | 5.169925 | 4.64385619 | 9.18239435 | 5.4918531 | 3.70043972 | 4.64385619 | 5.45943162 | 6.357552 | 4.32192809 | 6.22881869 | 4.24792751 | 4.64385619 |
|  |  |  | ZNF471 | 3.9068906 | 4.5849625 | 5.55458885 | 5 | 5 | 3.70043972 | 5.80735492 | 5.08746284 | 3.45943162 | 6.82017896 | 5.93073734 | 4.08746284 | 4.24792751 | 3.70043972 | 4.08746284 | 4.39231742 | 4.39231742 | 4.857981 | 4.80735492 |
|  |  |  | WNK3 | 2 | 1 | 2.5849625 | 2 | 1.5849625 | 1 | 2.32192809 | 2.32192809 | 1.5849625 | 0 | 6.71424552 | 2.80735492 | 2.32192809 | 0 | 1 | 1.5849625 | 0 | 0 | 1.5849625 |
|  |  |  | ZFP28 | 0 | 0 | 0 | 1 | 0 | 1 | 0 | 0 | 0 | 7.56224242 | 8.15987134 | 0 | 0 | 0 | 2.5849625 | 0 | 0 | 0 | 0 |
|  |  |  | HOXA4 | 0 | 5 | 6.32192809 | 6.10852446 | 4.52356196 | 0 | 4.64385619 | 4.5849625 | 3 | 2.32192809 | 8.4429435 | 0 | 1 | 0 | 2.5849625 | 5.97727992 | 5.04439412 | 6.94251451 | 5.7548875 |
|  |  |  | ZNF521 | 0 | 0 | 1.5849625 | 0 | 0 | 4.169925 | 5.61470984 | 0 | 3.169925 | 8.83605036 | 1 | 0 | 0 | 3.169925 | 3.32192809 | 0 | 7.48381578 | 0 | 0 |
|  |  |  | SYT15 | 4.857981 | 6.61470984 | 5.32192809 | 6.47573343 | 6.80735492 | 1.5849625 | 7.82017896 | 8.03891899 | 6.53915881 | 6.67242534 | 6.39231742 | 5.169925 | 5.80735492 | 7.86418614 | 5.7548875 | 6.22881869 | 6.87036472 | 6.42626475 | 5.95419631 |
|  |  |  | DOK6 | 0 | 1 | 0 | 0 | 0 | 0 | 1 | 0 | 0 | 6.79441587 | 6.93073734 | 0 | 0 | 0 | 2.5849625 | 0 | 0 | 0 | 0 |
|  |  |  | VGLL3 | 0 | 0 | 0 | 0 | 0 | 2 | 2.32192809 | 0 | 0 | 9.40301202 | 6.39231742 | 0 | 0 | 0 | 13.2206806 | 1 | 13.9775483 | 0 | 7 |
|  |  |  | PLXNA4 | 1 | 0 | 0 | 2.80735492 | 4.45943162 | 0 | 4.70043972 | 1 | 0 | 2 | 7.24792751 | 0 | 2 | 0 | 2 | 1 | 1.5849625 | 4.7548875 | 1 |
|  |  |  | USP51 | 0 | 0 | 3.70043972 | 0 | 0 | 2.5849625 | 1 | 0 | 3.169925 | 6.357552 | 4.70043972 | 0 | 0 | 2.80735492 | 3.45943162 | 0 | 2 | 0 | 5.39231742 |
|  |  |  | PCDHGA3 | 0 | 0 | 0 | 1.5849625 | 2.32192809 | 0 | 0 | 1 | 0 | 4.08746284 | 0 | 0 | 1.5849625 | 0 | 1.5849625 | 0 | 2.80735492 | 1.5849625 | 1.5849625 |
|  |  |  | MAGEL2 | 0 | 0 | 0 | 0 | 0 | 0 | 0 | 0 | 0 | 7.97154355 | 0 | 0 | 0 | 0 | 0 | 0 | 0 | 0 | 0 |

**Supplementary Table 8. The comparison of msEMT gene expression by the origin of gastric cancer cell lines.**

| Symbol | Ascites_Mean_exp. | Primary_Mean_exp. | Hemato_C_Mean_exp. | P_value  (Ascites vs. Primary) | P_value  (Ascites vs. Hemato) | P_value  (Primary vs. Hemato) | log2FC_Ascites vs. Primary) | log2FC_(Ascites vs. Hemato) | log2FC_(Primary vs. Hemato) |
| --- | --- | --- | --- | --- | --- | --- | --- | --- | --- |
| FMO1 | 0.2 | 0.264160417 | 0.861654167 | 0.850161829 | 0.286708175 | 0.335569965 | -0.064160417 | -0.661654167 | -0.59749375 |
| PREX2 | 0.6169925 | 2.306665178 | 0.528320834 | 0.313935879 | 0.887410898 | 0.30592197 | -1.689672678 | 0.088671667 | 1.778344345 |
| CDON | 7.098093533 | 6.970304794 | 6.607844022 | 0.892464988 | 0.554671419 | 0.74529568 | 0.127788739 | 0.490249511 | 0.362460772 |
| MPPED2 | 1.573216743 | 1.581975516 | 1 | 0.994304528 | 0.573138648 | 0.603604014 | -0.008758773 | 0.573216743 | 0.581975516 |
| PYGM | 2.876762524 | 3.369394492 | 3.806126573 | 0.43755073 | 0.210273248 | 0.56356526 | -0.492631968 | -0.929364049 | -0.436732082 |
| ABCC9 | 8.088082815 | 7.962621213 | 9.940525934 | 0.805874871 | 0.055342711 | 0.033582625 | 0.125461601 | -1.852443119 | -1.977904721 |
| RUNX1T1 | 0 | 1.497165596 | 0.935784974 | 0.126035031 | 0.422649731 | 0.669997689 | -1.497165596 | -0.935784974 | 0.561380622 |
| PCDHB4 | 1.287036472 | 1.894387036 | 2.803130312 | 0.418622702 | 0.124198829 | 0.353616595 | -0.607350564 | -1.51609384 | -0.908743276 |
| SMARCD3 | 7.582321642 | 8.30773909 | 5.403024179 | 0.505613699 | 0.281911723 | 0.183888903 | -0.725417448 | 2.179297463 | 2.904714912 |
| ZFHX4 | 3.278496921 | 3.07903549 | 1 | 0.919644575 | 0.026422334 | 0.277970637 | 0.199461431 | 2.278496921 | 2.07903549 |
| NFATC4 | 4.788608798 | 5.005819897 | 6.39868165 | 0.889018171 | 0.28745564 | 0.442291954 | -0.217211099 | -1.610072852 | -1.392861753 |
| KATNAL1 | 6.721361772 | 7.858296348 | 7.29425365 | 0.363749162 | 0.74841666 | 0.762053077 | -1.136934576 | -0.572891878 | 0.564042698 |
| CORO2B | 1.14918531 | 2.456893608 | 2.584962501 | 0.466891745 | 0.443695398 | 0.955086935 | -1.307708298 | -1.435777191 | -0.128068893 |
| IGDCC4 | 3.608087589 | 3.774128612 | 6.395872625 | 0.894879245 | 0.005730416 | 0.040095477 | -0.166041023 | -2.787785036 | -2.621744013 |
| NOVA2 | 2.88499548 | 1.652646563 | 2.566813239 | 0.199217985 | 0.831437778 | 0.58081635 | 1.232348917 | 0.318182241 | -0.914166676 |
| HOXA5 | 5.497053411 | 4.078393001 | 5.343785532 | 0.426930787 | 0.903422617 | 0.450253975 | 1.418660411 | 0.153267879 | -1.265392532 |
| GLI3 | 2.831058505 | 4.752535638 | 3.518515924 | 0.419742963 | 0.818117752 | 0.709133228 | -1.921477133 | -0.687457419 | 1.234019714 |
| EBF3 | 0.332192809 | 2.346135933 | 1.528320834 | 0.142872191 | 0.516976059 | 0.689406731 | -2.013943123 | -1.196128024 | 0.817815099 |
| TBC1D9 | 6.587625084 | 5.154990733 | 3.274455747 | 0.480701606 | 0.231780141 | 0.498598724 | 1.432634351 | 3.313169337 | 1.880534986 |
| NRXN2 | 2.730241357 | 2.465176852 | 3.056641667 | 0.775195153 | 0.740686056 | 0.513051381 | 0.265064505 | -0.32640031 | -0.591464815 |
| GLI1 | 4.388985934 | 5.614060441 | 2.555111972 | 0.3423076 | 0.327821274 | 0.148689749 | -1.225074506 | 1.833873962 | 3.058948469 |
| PHACTR1 | 1.460964047 | 3.037336754 | 2.659093308 | 0.012541347 | 0.163717807 | 0.613372329 | -1.576372706 | -1.19812926 | 0.378243446 |
| MDGA1 | 4.0244288 | 3.281466114 | 4.397430938 | 0.598368649 | 0.723138563 | 0.398759213 | 0.742962686 | -0.373002138 | -1.115964825 |
| EPM2A | 5.883176997 | 6.149883281 | 5.038943759 | 0.595030149 | 0.385345244 | 0.282378093 | -0.266706283 | 0.844233238 | 1.110939522 |
| PCDHB5 | 2.61262104 | 4.798737065 | 1.389975 | 0.345487934 | 0.239823652 | 0.14112347 | -2.186116025 | 1.222646039 | 3.408762065 |
| PDE8B | 5.921660215 | 4.918218094 | 4.886423747 | 0.24791355 | 0.236830182 | 0.943087311 | 1.003442121 | 1.035236468 | 0.031794346 |
| RASGRF2 | 3.154093054 | 4.48511566 | 2.854080808 | 0.288961639 | 0.547370481 | 0.188502941 | -1.331022606 | 0.300012246 | 1.631034852 |
| BCHE | 2.655937779 | 1.1949875 | 0 | 0.194364392 | 0.024795138 | 0.026836259 | 1.460950279 | 2.655937779 | 1.1949875 |
| PRRX1 | 0.445943162 | 1.881285397 | 0 | 0.301535965 | 0.23363388 | 0.182964229 | -1.435342235 | 0.445943162 | 1.881285397 |
| MYCT1 | 0.54918531 | 1.389975 | 0.666666667 | 0.205020209 | 0.882412002 | 0.439403794 | -0.840789691 | -0.117481357 | 0.723308334 |
| TNN | 0 | 1.385480493 | 0.666666667 | 0.299251601 | 0.183503419 | 0.58487289 | -1.385480493 | -0.666666667 | 0.718813826 |
| SMAD9 | 3.739743848 | 3.986395475 | 3.224141781 | 0.888682715 | 0.718382536 | 0.680774565 | -0.246651627 | 0.515602067 | 0.762253694 |
| TSHZ3 | 0.424792751 | 1.575201852 | 3.815866619 | 0.208984277 | 0.213451162 | 0.363843873 | -1.150409101 | -3.391073868 | -2.240664767 |
| NUDT10 | 0.2 | 2.52063975 | 0 | 0.232413755 | 0.167850656 | 0.199579064 | -2.32063975 | 0.2 | 2.52063975 |
| PDE1B | 0.89068906 | 1.317815099 | 2.699344028 | 0.553588063 | 0.501935313 | 0.601230383 | -0.42712604 | -1.808654968 | -1.381528928 |
| PLP1 | 0.25849625 | 1.329546654 | 0.666666667 | 0.209997867 | 0.613133698 | 0.524887174 | -1.071050404 | -0.408170417 | 0.662879987 |
| FAM124B | 0.4 | 0.861654167 | 0.666666667 | 0.312388496 | 0.523825149 | 0.716272633 | -0.461654167 | -0.266666667 | 0.1949875 |
| CCDC136 | 4.404593922 | 4.060504064 | 2.702841486 | 0.798787174 | 0.210965701 | 0.371217923 | 0.344089858 | 1.701752436 | 1.357662579 |
| ARHGAP22 | 4.855476536 | 4.589306217 | 4.917669256 | 0.561611684 | 0.915160381 | 0.637464476 | 0.266170319 | -0.062192721 | -0.328363039 |
| ATP1B2 | 2.341276416 | 2.580635963 | 2.535776771 | 0.87822786 | 0.858832048 | 0.975313574 | -0.239359547 | -0.194500355 | 0.044859192 |
| MAP7D3 | 8.284174117 | 8.615793831 | 5.74535987 | 0.743037244 | 0.179106488 | 0.142453693 | -0.331619714 | 2.538814247 | 2.870433961 |
| EDA2R | 1.467771964 | 1.496213321 | 3.099736006 | 0.974726484 | 0.483542592 | 0.487890726 | -0.028441356 | -1.631964042 | -1.603522686 |
| DLG4 | 5.723005028 | 6.779636371 | 7.275831557 | 0.388505746 | 0.340915433 | 0.758921095 | -1.056631343 | -1.552826529 | -0.496195186 |
| FHOD3 | 4.266374547 | 4.338362795 | 3.797439141 | 0.965559581 | 0.732181268 | 0.743052886 | -0.071988248 | 0.468935407 | 0.540923655 |
| ANKRD6 | 3.549753863 | 3.664618021 | 5.98047149 | 0.944701707 | 0.082910994 | 0.191306089 | -0.114864157 | -2.430717627 | -2.31585347 |
| KIAA1614 | 5.041939199 | 4.552288477 | 4.711903095 | 0.637388074 | 0.72916286 | 0.895760003 | 0.489650722 | 0.330036103 | -0.159614619 |
| NACAD | 3.817505549 | 3.227015608 | 3.15177574 | 0.740866317 | 0.677923904 | 0.970426415 | 0.590489941 | 0.665729809 | 0.075239868 |
| ARHGAP20 | 0.564385619 | 1.763219323 | 0.333333333 | 0.322205203 | 0.694960756 | 0.23392041 | -1.198833704 | 0.231052286 | 1.42988599 |
| TTLL7 | 5.480603754 | 5.781439792 | 4.36066089 | 0.763608797 | 0.131666299 | 0.140182979 | -0.300836038 | 1.119942864 | 1.420778902 |
| PRICKLE1 | 2.719589856 | 3.303298994 | 5.268295757 | 0.771091275 | 0.442106432 | 0.568569968 | -0.583709137 | -2.5487059 | -1.964996763 |
| PPFIA2 | 0.45849625 | 1.829546654 | 0 | 0.105771602 | 0.124179246 | 0.043367872 | -1.371050404 | 0.45849625 | 1.829546654 |
| MYOCD | 1.540354468 | 0.764160417 | 0 | 0.552718769 | 0.221273886 | 0.179003509 | 0.776194051 | 1.540354468 | 0.764160417 |
| RNF165 | 3.051998579 | 3.321831896 | 3.455440738 | 0.756382841 | 0.75844469 | 0.912021877 | -0.269833317 | -0.403442159 | -0.133608842 |
| CASQ1 | 1.364024494 | 1.640391724 | 0.333333333 | 0.783432998 | 0.100877259 | 0.203157782 | -0.27636723 | 1.03069116 | 1.307058391 |
| ANKRD53 | 0.94918531 | 1.041321252 | 1.861654167 | 0.901461231 | 0.402194316 | 0.49009389 | -0.092135943 | -0.912468857 | -0.820332915 |
| CAND2 | 3.124659383 | 4.021222194 | 3.071439707 | 0.585449354 | 0.914306975 | 0.557952166 | -0.896562811 | 0.053219676 | 0.949782487 |
| ZNF660 | 2.258836237 | 3.353939409 | 2.1949875 | 0.265356033 | 0.931011218 | 0.302775314 | -1.095103172 | 0.063848737 | 1.158951909 |
| ADPRH | 3.28908381 | 2.085949973 | 6.891062186 | 0.457083337 | 0.001607599 | 0.014219181 | 1.203133837 | -3.601978376 | -4.805112213 |
| NLGN4X | 0 | 3.601759833 | 0.333333333 | 0.118756185 | 0.422649731 | 0.15023001 | -3.601759833 | -0.333333333 | 3.2684265 |
| FAM13C | 0.778135971 | 1.166666667 | 0.528320834 | 0.684441756 | 0.719960816 | 0.538290156 | -0.388530695 | 0.249815138 | 0.638345833 |
| LRRC4C | 0.893663794 | 1.415308849 | 0 | 0.592998727 | 0.138510347 | 0.12480808 | -0.521645055 | 0.893663794 | 1.415308849 |
| KIRREL3 | 2.620609861 | 3.771257725 | 3.678724538 | 0.344272632 | 0.223653311 | 0.940525816 | -1.150647864 | -1.058114676 | 0.092533188 |
| KIAA1755 | 1.04918531 | 1.946286607 | 3.224730756 | 0.151096284 | 0.316210948 | 0.522969342 | -0.897101297 | -2.175545446 | -1.278444149 |
| CNKSR2 | 1.004439412 | 2.36414589 | 0.333333333 | 0.390595353 | 0.326927583 | 0.203973641 | -1.359706479 | 0.671106079 | 2.030812557 |
| MKX | 1.42467406 | 1.560323962 | 2.344474334 | 0.906641489 | 0.735056179 | 0.780364137 | -0.135649902 | -0.919800274 | -0.784150372 |
| FAM124A | 3.621110765 | 4.395987237 | 2.60459373 | 0.403075401 | 0.453212936 | 0.214865029 | -0.774876472 | 1.016517035 | 1.791393507 |
| GPM6A | 0.970043972 | 2.9340729 | 0.666666667 | 0.21313549 | 0.734944784 | 0.171997001 | -1.964028928 | 0.303377305 | 2.267406234 |
| DLG2 | 3.789350701 | 3.499941289 | 4.38893025 | 0.663766745 | 0.29235163 | 0.064203767 | 0.289409411 | -0.599579549 | -0.888988961 |
| DIXDC1 | 7.035256197 | 7.856239856 | 6.482609954 | 0.210234308 | 0.496800295 | 0.135362625 | -0.820983659 | 0.552646243 | 1.373629902 |
| AKAP6 | 5.195851166 | 4.763509595 | 5.032896699 | 0.667729342 | 0.78120757 | 0.765217292 | 0.432341571 | 0.162954468 | -0.269387104 |
| CACNA2D1 | 3.309970352 | 4.357556998 | 1.528320834 | 0.631170402 | 0.394658378 | 0.271782737 | -1.047586646 | 1.781649518 | 2.829236164 |
| FGD5 | 3.560200565 | 3.961390173 | 3.664784479 | 0.379109262 | 0.799777793 | 0.584438586 | -0.401189608 | -0.104583914 | 0.296605694 |
| KCNMA1 | 2.537054268 | 4.668623271 | 1.389975 | 0.354599439 | 0.374893218 | 0.162821653 | -2.131569003 | 1.147079268 | 3.27864827 |
| ADAMTSL3 | 6.901066155 | 6.6992735 | 8.539029655 | 0.656971612 | 0.010499616 | 0.007983504 | 0.201792655 | -1.637963499 | -1.839756154 |
| ERG | 0.696578428 | 0.386988016 | 0 | 0.630107339 | 0.19342206 | 0.363217468 | 0.309590413 | 0.696578428 | 0.386988016 |
| SHROOM4 | 2.230563429 | 3.335029734 | 2.497284365 | 0.106458494 | 0.858060792 | 0.586563958 | -1.104466305 | -0.266720937 | 0.837745369 |
| DYNC1I1 | 2.319315956 | 4.942377568 | 6.661202001 | 0.086267707 | 8.44E-05 | 0.205278659 | -2.623061612 | -4.341886045 | -1.718824433 |
| WNT9B | 1.733985 | 2.184427157 | 0.333333333 | 0.347845332 | 0.023324796 | 0.008405356 | -0.450442156 | 1.400651667 | 1.851093823 |
| C1QTNF7 | 0.15849625 | 0 | 0 | 0.343436396 | 0.343436396 | NA | 0.15849625 | 0.15849625 | 0 |
| TDRD10 | 0.4169925 | 0.940642698 | 0.666666667 | 0.340406825 | 0.750318315 | 0.752798168 | -0.523650198 | -0.249674167 | 0.273976032 |
| MAP9 | 2.891242469 | 2.805292096 | 2.396608186 | 0.969128333 | 0.865570555 | 0.897275462 | 0.085950373 | 0.494634283 | 0.408683909 |
| CMYA5 | 3.864545337 | 3.468627383 | 6.12324455 | 0.560214371 | 0.000285139 | 0.003113747 | 0.395917954 | -2.258699213 | -2.654617167 |
| MYOZ3 | 1.54918531 | 3.112305589 | 1.635630199 | 0.283359783 | 0.884933881 | 0.305656126 | -1.56312028 | -0.086444889 | 1.476675391 |
| PKNOX2 | 0.4169925 | 2.088401056 | 1.153143873 | 0.195649321 | 0.590727247 | 0.582129533 | -1.671408556 | -0.736151373 | 0.935257183 |
| TTC7B | 6.08077093 | 5.986007769 | 7.686138197 | 0.965452534 | 0.134200385 | 0.423331359 | 0.094763161 | -1.605367267 | -1.700130428 |
| YPEL4 | 0.933985 | 1.6949875 | 1.602451641 | 0.318421894 | 0.526922065 | 0.931675946 | -0.7610025 | -0.66846664 | 0.09253586 |
| MAP1A | 5.229218701 | 5.719311185 | 5.608895008 | 0.749606079 | 0.740466538 | 0.941313636 | -0.490092484 | -0.379676307 | 0.110416177 |
| GFRA2 | 0.45849625 | 1.229173239 | 0.333333333 | 0.567618353 | 0.796149081 | 0.509563362 | -0.770676988 | 0.125162917 | 0.895839905 |
| FAM110B | 3.424263827 | 4.406544385 | 0 | 0.641958128 | 0.015013852 | 0.048423453 | -0.982280558 | 3.424263827 | 4.406544385 |
| CPT1C | 5.005561566 | 6.368738141 | 5.157309729 | 0.463750302 | 0.919926753 | 0.56247879 | -1.363176575 | -0.151748163 | 1.211428412 |
| MN1 | 3.605346667 | 3.489093177 | 3.163951032 | 0.952570859 | 0.769553421 | 0.858381141 | 0.116253491 | 0.441395635 | 0.325142145 |
| ZFPM2 | 2.989059679 | 5.351895464 | 4.703004806 | 0.050165927 | 0.002733356 | 0.519962787 | -2.362835786 | -1.713945128 | 0.648890658 |
| LRRN2 | 3.707915112 | 2.854625427 | 3.082642504 | 0.567527374 | 0.775039247 | 0.920798369 | 0.853289685 | 0.625272608 | -0.228017077 |
| DSEL | 3.486736137 | 3.738005153 | 3.233479906 | 0.885349494 | 0.829007684 | 0.71481803 | -0.251269016 | 0.253256231 | 0.504525247 |
| ZNF483 | 3.764304016 | 4.43691853 | 3.478903847 | 0.447374501 | 0.701312032 | 0.310078702 | -0.672614514 | 0.285400169 | 0.958014683 |
| ZNF135 | 0.1 | 0.166666667 | 0 | 0.739809977 | 0.343436396 | 0.363217468 | -0.066666667 | 0.1 | 0.166666667 |
| CCDC89 | 1.86707254 | 1.151148433 | 0.861654167 | 0.395090861 | 0.278712109 | 0.632369033 | 0.715924107 | 1.005418373 | 0.289494266 |
| SSC5D | 2.139196516 | 3.925706098 | 3.39330303 | 0.053977382 | 0.264320195 | 0.643178474 | -1.786509582 | -1.254106514 | 0.532403068 |
| TMEM132E | 0.75849625 | 2.095962597 | 0 | 0.269872507 | 0.112664914 | 0.096665136 | -1.337466347 | 0.75849625 | 2.095962597 |
| GPR135 | 4.528966503 | 4.298748708 | 2.974937501 | 0.728633188 | 0.004230906 | 0.052456731 | 0.230217795 | 1.554029001 | 1.323811206 |
| UNC5C | 1.316741815 | 1.475291507 | 0.333333333 | 0.888134906 | 0.236660706 | 0.251960007 | -0.158549692 | 0.983408481 | 1.141958174 |
| HHIPL1 | 2.178676098 | 2.80862484 | 1 | 0.669495152 | 0.241187313 | 0.22424252 | -0.629948743 | 1.178676098 | 1.80862484 |
| TRIM61 | 0.9 | 0.166666667 | 1.528320834 | 0.08081423 | 0.202227702 | 0.02113277 | 0.733333333 | -0.628320834 | -1.361654167 |
| FBXL7 | 1 | 3.410566108 | 0 | 0.188987522 | 0.084785213 | 0.07873885 | -2.410566108 | 1 | 3.410566108 |
| PCDH9 | 2.31944587 | 2.953685627 | 0 | 0.741251098 | 0.043164269 | 0.120342683 | -0.634239757 | 2.31944587 | 2.953685627 |
| PRKD1 | 1.56617781 | 3.454667392 | 2.02915428 | 0.301427141 | 0.84291001 | 0.607110523 | -1.888489583 | -0.462976471 | 1.425513112 |
| SORCS2 | 2.869479316 | 1.645060787 | 0.666666667 | 0.288068953 | 0.058757422 | 0.147306421 | 1.224418529 | 2.202812649 | 0.97839412 |
| C3orf70 | 2.702810031 | 1.776963016 | 2.464105808 | 0.257555548 | 0.560646833 | 0.379631741 | 0.925847015 | 0.238704224 | -0.687142791 |
| MAGI2 | 3.490935335 | 1.93300875 | 4.893526432 | 0.310880643 | 0.269604504 | 0.100929665 | 1.557926585 | -1.402591097 | -2.960517682 |
| GREB1 | 4.965875344 | 5.506077544 | 5.447867379 | 0.541037761 | 0.320235918 | 0.948196512 | -0.540202201 | -0.481992036 | 0.058210165 |
| ZNF471 | 4.444773279 | 4.97300448 | 5.046517117 | 0.363722614 | 0.154458322 | 0.901320872 | -0.5282312 | -0.601743838 | -0.073512638 |
| WNK3 | 1.320609861 | 2.270189352 | 1.861654167 | 0.38005785 | 0.391270212 | 0.710956571 | -0.949579491 | -0.541044306 | 0.408535185 |
| ZFP28 | 0.25849625 | 2.78701896 | 0.333333333 | 0.179756074 | 0.866501812 | 0.192390435 | -2.52852271 | -0.074837083 | 2.453685627 |
| HOXA4 | 3.349417677 | 3.554519002 | 5.810150851 | 0.900192853 | 0.016737391 | 0.171260606 | -0.205101325 | -2.460733173 | -2.255631848 |
| ZNF521 | 2.276030372 | 2.334329226 | 0.528320834 | 0.973333341 | 0.113325962 | 0.287454705 | -0.058298854 | 1.747709538 | 1.806008393 |
| SYT15 | 6.501387205 | 5.629237323 | 6.137457123 | 0.394514632 | 0.502345013 | 0.626940146 | 0.872149882 | 0.363930082 | -0.5082198 |
| DOK6 | 0.35849625 | 2.287525534 | 0.333333333 | 0.243331634 | 0.955305261 | 0.240251498 | -1.929029284 | 0.025162917 | 1.954192201 |
| VGLL3 | 3.652015697 | 3.132554908 | 0 | 0.831688659 | 0.072890923 | 0.10512413 | 0.51946079 | 3.652015697 | 3.132554908 |
| PLXNA4 | 2.049972134 | 2.041321252 | 0.935784974 | 0.994583788 | 0.376923248 | 0.465414804 | 0.008650882 | 1.11418716 | 1.105536278 |
| USP51 | 1.782902896 | 2.273825704 | 1.233479906 | 0.710791782 | 0.715289652 | 0.55961905 | -0.490922807 | 0.54942299 | 1.040345798 |
| PCDHGA3 | 1.146913302 | 0.847910474 | 0.528320834 | 0.700354814 | 0.381742837 | 0.719148145 | 0.299002828 | 0.618592468 | 0.31958964 |
| MAGEL2 | 0 | 1.328590592 | 0 | 0.363217468 | NA | 0.363217468 | -1.328590592 | 0 | 1.328590592 |
